# Protection against SARS-CoV-2 Beta Variant in mRNA-1273 Boosted Nonhuman Primates

**DOI:** 10.1101/2021.08.11.456015

**Authors:** Kizzmekia S. Corbett, Matthew Gagne, Danielle A. Wagner, Sarah O’ Connell, Sandeep R. Narpala, Dillon R. Flebbe, Shayne F. Andrew, Rachel L. Davis, Barbara Flynn, Timothy S. Johnston, Christopher Stringham, Lilin Lai, Daniel Valentin, Alex Van Ry, Zackery Flinchbaugh, Anne P. Werner, Juan I. Moliva, Manjari Sriparna, Sijy O’Dell, Stephen D. Schmidt, Courtney Tucker, Angela Choi, Matthew Koch, Kevin W. Bock, Mahnaz Minai, Bianca M. Nagata, Gabriela S. Alvarado, Amy R. Henry, Farida Laboune, Chaim A. Schramm, Yi Zhang, Lingshu Wang, Misook Choe, Seyhan Boyoglu-Barnum, Wei Shi, Evan Lamb, Saule T. Nurmukhambetova, Samantha J. Provost, Mitzi M. Donaldson, Josue Marquez, John-Paul M. Todd, Anthony Cook, Alan Dodson, Andrew Pekosz, Eli Boritz, Aurélie Ploquin, Nicole Doria-Rose, Laurent Pessaint, Hanne Andersen, Kathryn E. Foulds, John Misasi, Kai Wu, Andrea Carfi, Martha C. Nason, John Mascola, Ian N. Moore, Darin K. Edwards, Mark G. Lewis, Mehul S. Suthar, Mario Roederer, Adrian McDermott, Daniel C. Douek, Nancy J. Sullivan, Barney S. Graham, Robert A. Seder

**Affiliations:** Vaccine Research Center; National Institute of Allergy and Infectious Diseases; National Institutes of Health; Bethesda, Maryland, 20892; United States of America; Center for Childhood Infections and Vaccines of Children’s Healthcare of Atlanta, Department of Pediatrics, Department of Microbiology and Immunology, Emory Vaccine Center, Emory University, Atlanta, Georgia, 30322, United States of America; Bioqual, Inc.; Rockville, Maryland, 20850; United States of America; Moderna Inc., Cambridge, MA, 02139; United States of America; Infectious Disease Pathogenesis Section; National Institute of Allergy and Infectious Diseases; National Institutes of Health; Bethesda, Maryland, 20892; United States of America; Department of Microbiology and Immunology; Johns Hopkins University, Bloomberg School of Public Health, Baltimore, Maryland, 21205, United States of America; Biostatistics Research Branch, Division of Clinical Research, National Institute of Allergy and Infectious Diseases, National Institutes of Health; Bethesda, Maryland, 20892; United States of America; Department of Microbiology and Immunology, Harvard T.H. Chan School of Public Health; Boston, Massachusetts, 02115; United States of America; Duke University School of Medicine; Medical Sciences Program; Durham, North Carolina, 27705; United States of America; Medical College of Virginia; Richmond, Virginia, 23223; United States of America

**Author notes:** These authors contributed equally to this manuscript (Kizzmekia S. Corbett, Matthew Gagne).

## Abstract

Neutralizing antibody responses gradually wane after vaccination with mRNA-1273 against several variants of concern (VOC), and additional boost vaccinations may be required to sustain immunity and protection. Here, we evaluated the immune responses in nonhuman primates that received 100 µg of mRNA-1273 vaccine at 0 and 4 weeks and were boosted at week 29 with mRNA-1273 (homologous) or mRNA-1273.β (heterologous), which encompasses the spike sequence of the B.1.351 (beta or β) variant. Reciprocal ID_50_ pseudovirus neutralizing antibody geometric mean titers (GMT) against live SARS-CoV-2 D614G and the β variant, were 4700 and 765, respectively, at week 6, the peak of primary response, and 644 and 553, respectively, at a 5-month post-vaccination memory time point. Two weeks following homologous or heterologous boost β-specific reciprocal ID_50_ GMT were 5000 and 3000, respectively. At week 38, animals were challenged in the upper and lower airway with the β variant. Two days post-challenge, viral replication was low to undetectable in both BAL and nasal swabs in most of the boosted animals. These data show that boosting with the homologous mRNA-1273 vaccine six months after primary immunization provides up to a 20-fold increase in neutralizing antibody responses across all VOC, which may be required to sustain high-level protection against severe disease, especially for at-risk populations.

**One-sentence summary:** mRNA-1273 boosted nonhuman primates have increased immune responses and are protected against SARS-CoV-2 beta infection.

## INTRODUCTION

SARS-CoV-2 vaccines, such as mRNA-1273, BNT162b2, AD26.COV2.S, AZD1222, and NVX-CoV2373, which deliver the Wuhan-1 spike (S) protein are between 70-94% effective against symptomatic D614G or B.1.1.7 alpha (α) variant infection (*1–5*). Several SARS-CoV-2 variants of concern (VOC) have reduced sensitivity to neutralizing antibodies induced by vaccination or prior infection. The B.1.617.2 delta (δ) variant is notable for its high transmissibility and an intermediate level of reduced neutralization by WA-1 immune sera (*6, 7*). The B.1.351 beta (β) variant - initially isolated in South Africa - and the P.1 gamma (*γ*) variant - initially isolated in Brazil - show the greatest reduction in neutralizability by vaccinee or convalescent patient sera (*8–13*).

Since mRNA-1273 was granted Emergency Use Authorization (EUA) in December 2020 and deployed globally, months have elapsed since the first subjects were vaccinated, and durability of protection remains a concern. Additionally, the question of a need for an additional booster immunization is clinically relevant, particularly as new VOC become more prevalent. mRNA-1273 vaccination in humans induced antibody responses that persisted beyond 6 months (*14*), but serum neutralizing activity was significantly reduced against β and δvariants (*7, 15, 16*). Recent data show that boosting subjects fully vaccinated with mRNA-1273 several months after initial priming with either mRNA-1273, mRNA-1273.β (S sequence based on the β variant), or a 1:1 combination of both vaccines (mRNA-1273.211) showed significantly increased neutralizing titers against the prototypic D614G virus and all variants, including β (*17*). While these *in vitro* functional data are promising, there are no available efficacy data to define the level of antibodies that are sufficient to mediate protection against infection since the correlates of protection against VOCs are undetermined. Moreover, in NHP, a higher threshold of neutralizing antibodies was needed for upper compared to lower airway protection (*18*), suggesting a boost could be important to reach this threshold and contribute to reducing virus transmission.

Nonhuman primates (NHP) have proven useful for assessing immunogenicity and protection against SARS-CoV-2 to inform vaccine development (*19–24*). Here, NHP immunized with the clinically relevant mRNA-1273 vaccine regimen (100 µg x2) were boosted ∼6 months later with mRNA-1273 (homologous) or mRNA-1273.β (heterologous). Antibody, B cell, and T cell responses were assessed temporally to determine how boosting with mRNA-1273 or mRNA-1273.β influences the magnitude of WA-1 or β-specific immune responses and protection against SARS-CoV-2 β challenge. The data show that an additional, delayed boost with mRNA vaccines matched to either the original vaccine strain or to the heterologous challenge strain can have a striking effect on increasing the breadth of neutralization against all VOC tested and protection against the βvariant. The data presented here provide a rationale for how boosting can influence immunity and protection in humans.

## RESULTS

### Boosting increases mRNA-1273-induced serum antibody responses

The primary focus of this study was to assess the effect of homologous or heterologous boosting with the original WA-1 mRNA-1273 or mRNA-1273.β, respectively, on immunogenicity and protection against infection with the β variant. NHP that received the clinically relevant primary regimen of mRNA-1273 (100 µg at weeks 0 and 4) were boosted with either mRNA-1273 (homologous) or mRNA-1273.β (heterologous) (**Fig. S1**). WA-1 and β S- or receptor binding domain (RBD)-specific serum antibody responses were assessed at weeks 6, 24 or 29, 31, and 37, which correspond to peak (*19*), memory pre-boost, 2 weeks post-boost, and time of challenge (“pre-challenge”) responses, respectively. WA-1 and β S-specific IgG geometric mean titers (GMT) were 53,000 and 39,000 AUC, respectively, at the peak timepoint following a primary regimen of mRNA-1273. These responses dropped 4- and 6-fold, respectively, after ∼6 months (week 29). Boosting with mRNA-1273 or mRNA-1273.β induced 4- and ∼6-fold increases in WA-1 and β S-specific IgG titers such that at 2 weeks following the memory boost, WA-1 and β S-specific IgG GMT were restored to peak levels **(Fig. 1A-B)**. By week 37, S-specific IgG GMT remained >30,000 and >20,000 AUC, respectively for WA-1 and β **(Fig. 1A-B)**, which, for WA-1, translated to >200,000 international units (IU/mL) **(Table S1)**. RBD-specific responses displayed similar kinetic trends **(Fig. 1C-D)**, and together these data show there is no difference in the ability of mRNA-1273 or mRNA-1273.β to boost primary mRNA-1273-elicited S-binding IgG responses.

**Figure 1.**
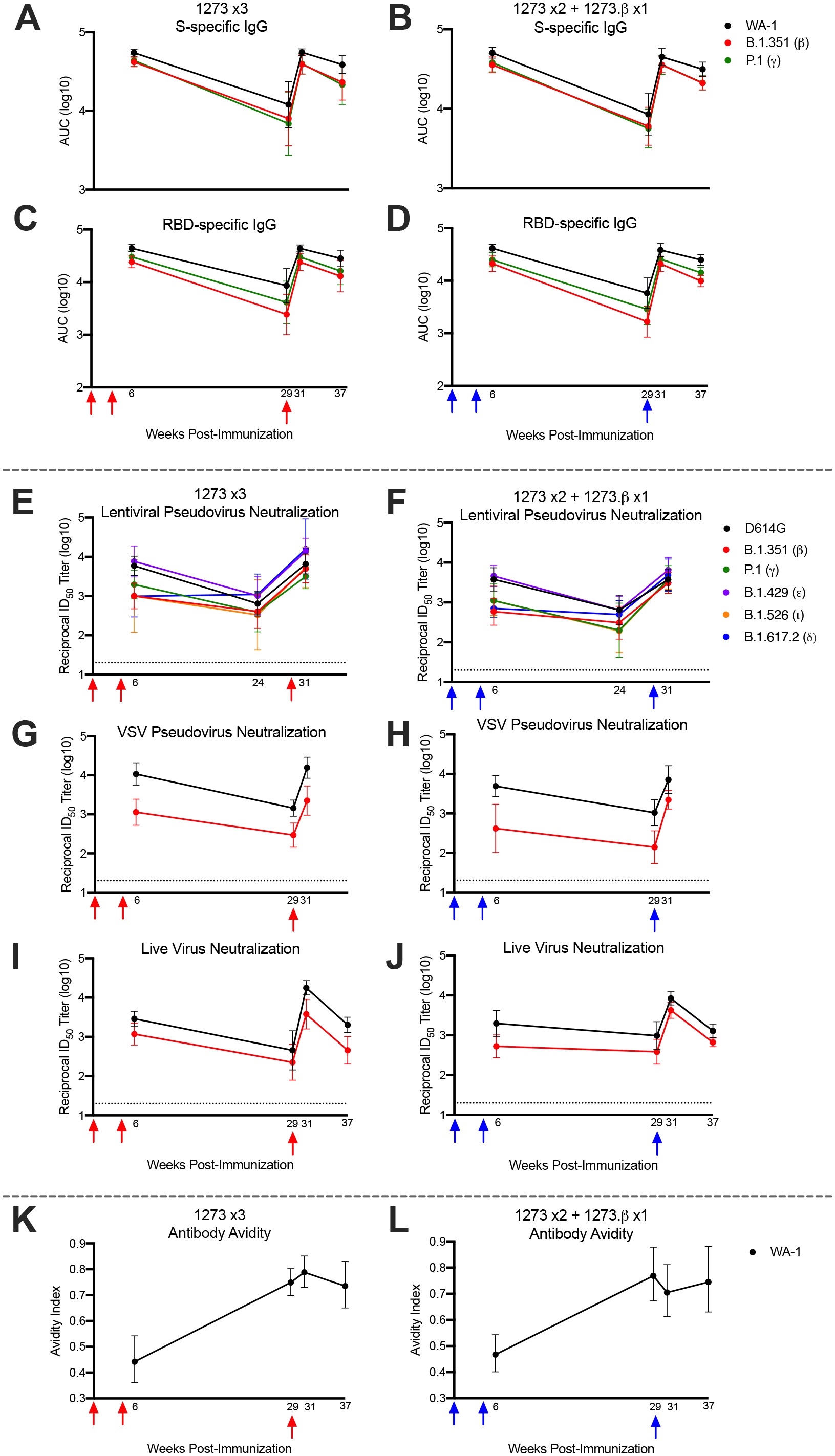
Temporal serum antibody responses to SARS-CoV-2 variants. Rhesus macaques (n = 8/group) were immunized according to Figure S1. Sera were collected at weeks 6, 24 or 29, 31, and 37 and assessed, where applicable, for SARS-CoV-2 WA-1 (black), β (red), and P.1 – gamma, *γ* (green), B.1.429 – epsilon, *ε* (purple), B.1.526 – iota, *ι* (orange), and B.1.617.2 – delta, δ (blue) S-specific (A-B) and RBD-specific (C-D) IgG by MULTI-ARRAY ELISA, lentiviral-based pseudovirus neutralization (E-F), VSV-based pseudovirus neutralization (G-H), live virus neutralization (I-J), and WA-1 S-specific antibody avidity (K-L). Lines represent geometric mean titer (GMT) and geometric error. Dotted lines indicate neutralization assay limits of detection. Arrows point to immunization timepoints.

Next, we assessed the kinetics of neutralizing antibody responses using a lentiviral-based neutralizing antibody assay for D614G, the benchmark strain, as well as several globally circulating variants (β, δ, P.1 - *γ*, B.1.429 - *ε*, and B.1.526 - *ι*). Consistent with previous data in NHP and humans (*7, 15*), mRNA-1273 elicited a variant-dependent hierarchy of neutralizing antibody responses; at peak, the ID_50_ GMT for D614G was 4700, followed by 5900, 1500, 1100, 834, and 765 for ε, γ, ι, δ, and β, respectively, representing 1-6-fold decreases compared to D614G. At the memory timepoint (week 24), neutralizing antibody titers decreased to 644 ID_50_ GMT for D614G and 808, 278, 252, 738, and 553 for ε, γ, ι, δ, and β, respectively. Of note, there was a greater fold reduction in neutralization titers from week 6 to week 24 for D614G compared to β and δ (*p* <0.0001) **(Fig. 1E-F)**. Similar observations were made using D614G and β VSV-based pseudovirus (**Fig. 1G-H**) or live virus neutralization assays (**Fig. 1I-J**). Last, WA-1 S-specific antibody avidity, one measure of affinity maturation, was significantly increased over the 6 months following the primary mRNA-1273 immunization series in these NHP (*p* <0.0001) **(Fig. 1K-L)**. This indicates continued affinity maturation occurs following primary immunization leading to an increase in antibody quality and suggests that, for some variants, affinity maturation of mRNA-1273-elicited neutralizing antibody responses may occur over time, even in the absence of continued antigen exposure.

Six months after the primary immunization series, either a homologous (mRNA-1273) or heterologous (mRNA-1273.β) boost induced a 7-21-fold increase in neutralizing antibody titers for all the variants two weeks later, resulting in, for example, 5000 and 3000 β-specific ID_50_ GMT following mRNA-1273 or mRNA-1273.βboosts, respectively **(Fig. 1E-F)**. Of note, these responses to variants were significantly higher than at week 6 for both boost groups (δ-specific neutralizing antibody responses, *p* = 0.002 and 0.003 for mRNA-1273 and mRNA-1273.β boosts, respectively). This post-boost increase in lentiviral pseudovirus neutralizing titers was confirmed using a VSV-based pseudovirus neutralization assay **(Fig. 1G-H)** and a live virus neutralization assay **(Fig. 1I-J)**. By week 37, live virus neutralizing antibody titers returned to week 6 levels **(Fig. 1I-J)**. Together, these data show that despite waning of mRNA-1273-elicited antibody responses over 6 months, boosting with either homologous or heterologous mRNA was able to restore antibody responses to peak levels and, importantly, with sufficient potency and breadth to neutralize across heterologous virus variants.

### Boosting increases antibody responses in the airway

Vaccine-induced antibodies localized in the upper and lower airways may play a role in the initial control of SARS-CoV-2 replication (*18, 25, 26*). Therefore, we extended analysis beyond circulating antibody responses to assess WA-1 and β S-specific IgG responses in bronchoalveolar lavage (BAL) and nasal wash samples (NS) at weeks 6 and 36. At week 6, two doses of mRNA-1273 elicited WA-1 and β S-specific GMTs of 13 and 11 AUC, respectively, in the BAL (**Figs. 2A and 2C**). Similar GMT (11 and 10 AUC) of S-specific IgG were also detected in nasal wash samples (**Figs. 2B and 2D**). Interestingly, following a boost with either mRNA-1273 or mRNA-1273.β, the levels of BAL and nasal wash S-specific IgG were also 10-12 AUC GMT **(Figs. 2A-D)**. To detect functional antibodies in these mucosal samples, we utilized an ACE2 binding inhibition assay. In the BAL of NHP boosted with mRNA-1273, ACE2 binding to WA-1 and β S proteins was reduced by a mean of 54% and 37%, respectively (**Fig. 2E**), and 47% and 32% in nasal washes **(Fig. 2F)**. Similarly, in BAL and nasal washes of NHP that were boosted with mRNA-1273.β, ACE2 binding to WA-1 S was reduced by 42% and 55% and for β S proteins by 30% and 46%, respectively; ACE2 binding inhibition was reduced for β as compared to WA-1, similar to what was observed for serum antibodies (**Figs. 2E-F**). These data confirm that either homologous or heterologous memory vaccination can induce upper and lower airway functional antibodies that are necessary to mitigate illness and transmission.

**Figure 2.**
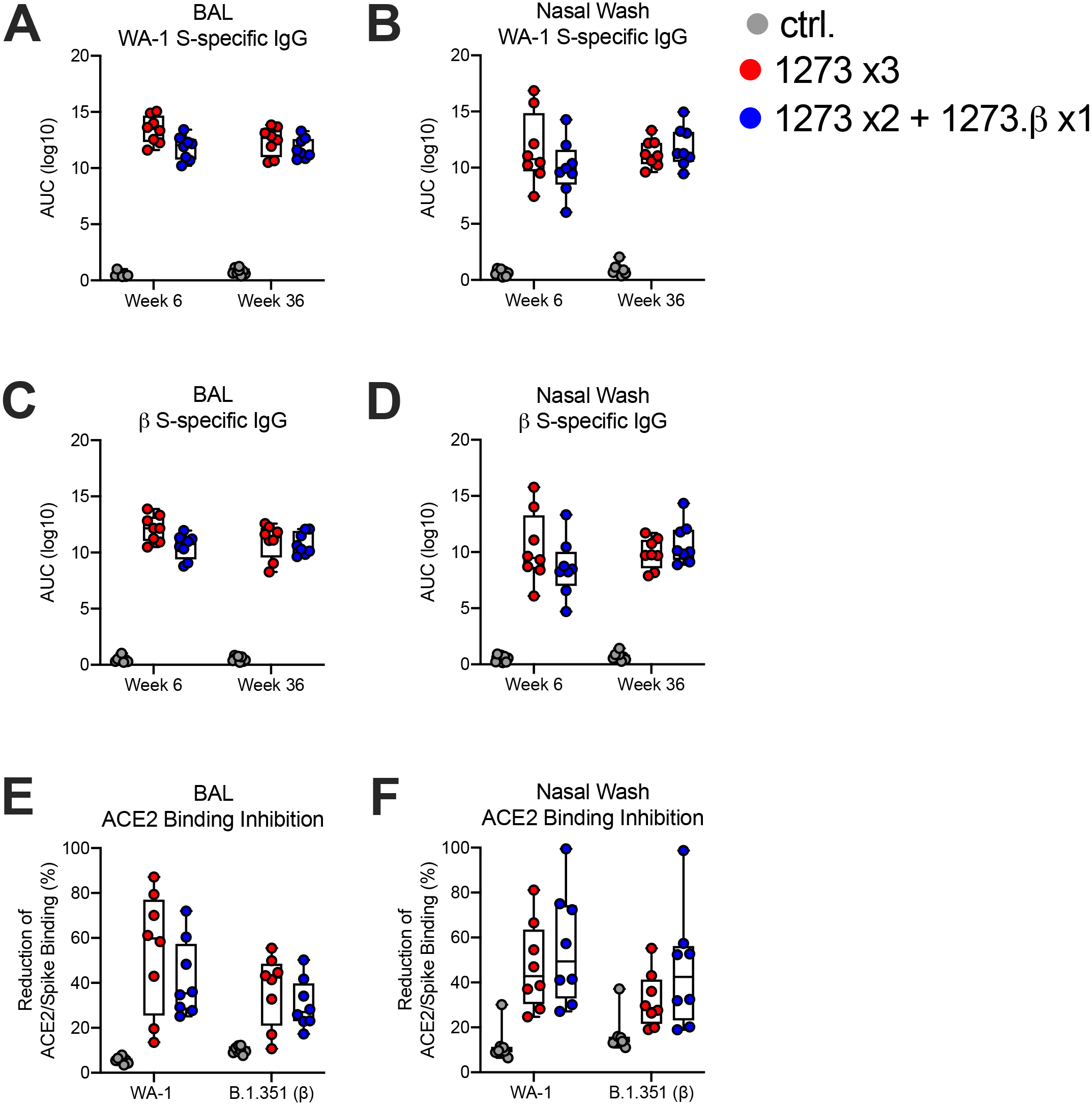
Mucosal antibody responses to SARS-CoV-2 variants. Rhesus macaques (n = 8/group) were immunized according to Figure S1. BAL (A, C, E) and nasal washes (B, D, F) collected at weeks 6 and 36 were assessed for S-specific IgG to SARS-CoV-2 WA-1 (A-B) and β (C-D) by MULTI-ARRAY ELISA. Week 36 BAL and nasal washes were assessed for inhibition of ACE2 binding to WA-1 and β S (E-F). Boxes and horizontal bars denote interquartile ranges (IQR) and medians, respectively; whisker end points are equal to the maximum and minimum values. Circles represent individual NHP.

### mRNA-1273.β immunization induces robust serum and mucosal antibody responses

A secondary focus of this study was to evaluate the immunogenicity of mRNA-1273.βas a primary vaccine regimen as this may have relevance for design of future vaccines in naïve individuals; this group additionally served as a homologous vaccine control for the β challenge **(Fig. S1)**. S-specific IgG GMT for WA-1 and β were 5100 and 8900 two weeks after the first immunization of mRNA-1273.β and increased 4-5-fold after a second immunization **(Fig. S2A)**. Two doses of mRNA-1273.β resulted in D614G and β live virus ID_50_ GMT of 198 and 788, respectively **(Fig. S2B)**. Potent S-specific antibody responses against WA-1 or β were also detected in BAL (**Fig. S2C**) and nasal washes (**Fig. S2D**). Mucosal antibody responses were further exhibited by ACE2 binding inhibition, where mRNA-1273.β induced BAL antibodies that reduced ACE2 binding to WA-1 and β S by a median of 21% and 48%, respectively (**Fig. 2E)**. In nasal washes, median reduction of ACE2 binding to WA-1 and β S binding was 30% and 56%, respectively (**Fig. 2F)**. Together, these data show mRNA-1273.β given as a primary regimen elicits higher β-specific responses as compared to WA-1.

### Serum antibody repertoire elicited by homologous or heterologous prime boost

To evaluate the impact of homologous (WA-1) or heterologous (β) boosting antigen on serum antibody epitope specificity, the absolute value (RUs) and the relative proportion (percent competition) of serum antibodies against 16 distinct antigenic sites (**Table S2**) on WA-1 SARS-CoV-2 S was measured using surface plasmon resonance (SPR). We evaluated serum antibody specificity at week 35 (6 weeks post-boost), to allow sufficient time for B cell expansion following exposure to naïve antigen in heterologous-boosted animals. The epitope specificity spanned the S2, N-terminal domain (NTD), S1, and RBD subdomains of S protein and the breadth was similar whether the boost was homologous or heterologous (**Fig 3A**). The specificity of serum antibodies was qualitatively similar across all animals within each group for the NTD subdomain (average standard deviation 32.1-36.9), but not for RBD subdomain (average standard deviation 50.2-56.1) (**Fig 3A**). Between vaccine groups, there was more NTD site A specificity in homologous than in heterologous boosted animals (**Fig. 3A**). Both absolute and relative serum reactivity to NTD site A (mAb 4-8, **Table S2**) were higher in homologous than heterologous boosted animals at all time points post-boost (**Fig. 3B**), indicating mutations found in the β S protein (Δ242-244, R246I) may render the heterologous boost unable to recall a primary mRNA-1273 immunization memory B-cell response to this antigenic site. In contrast, RBD site H (mAb A23-97.1, **Table S2**) showed differences in relative but not absolute serum reactivity, with heterologous inducing higher reactivity than homologous boosting at all post memory boost time points (**Fig 3C**). Because A23-97.1 binds equivalently to both WA-1 and β S (**Fig. S3**), the higher relative (but not absolute) reactivity against this site for β is likely due to the decreased contribution of NTD-A reactivity, as shown by the ratio of mRNA-1273.β to mRNA-1273 relative reactivity **(Fig. 3D)**. Overall, mRNA-1273 to β ratios decrease over time, indicating a contraction of the naïve β-directed response and return to a memory, long-lived epitope profile (**Fig. 3D**), and suggesting heterologous boost does not alter absolute epitope reactivity over time. In contrast, following primary immunization with mRNA-1273.β we observed significantly increased absolute serum reactivity to RBD sites B, C, D, and F (**Fig. S4**) as compared to homologous boost, with sites C and D (A19-46.1, A19-61.1, **Table S2**) being associated with strong neutralization potency against β (*27*). These data highlight differences between primary immunization and heterologous boost with mRNA-1273.β and indicate primary immunization plays a key role in shaping the overall serum antibody epitope repertoire.

**Figure 3.**
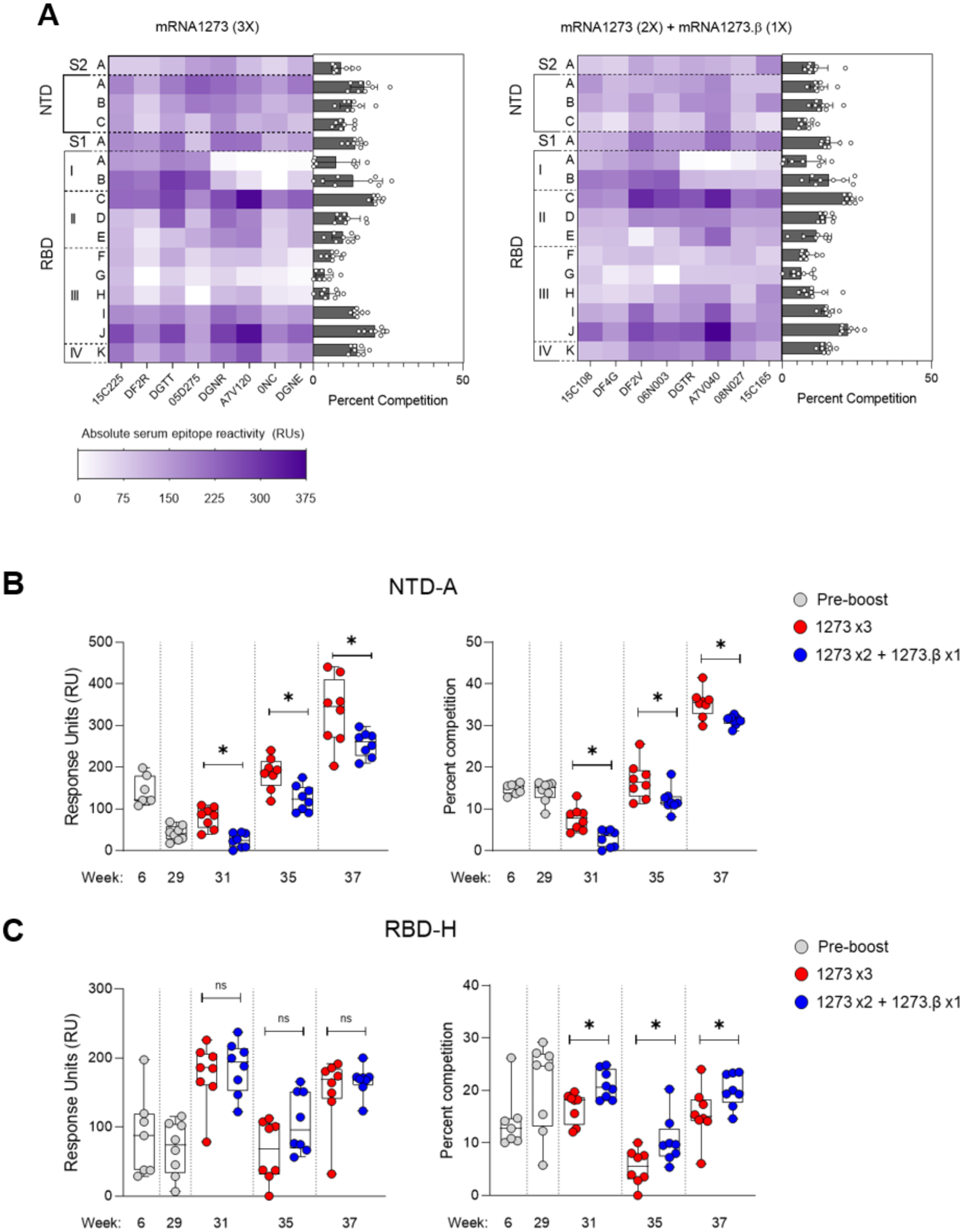
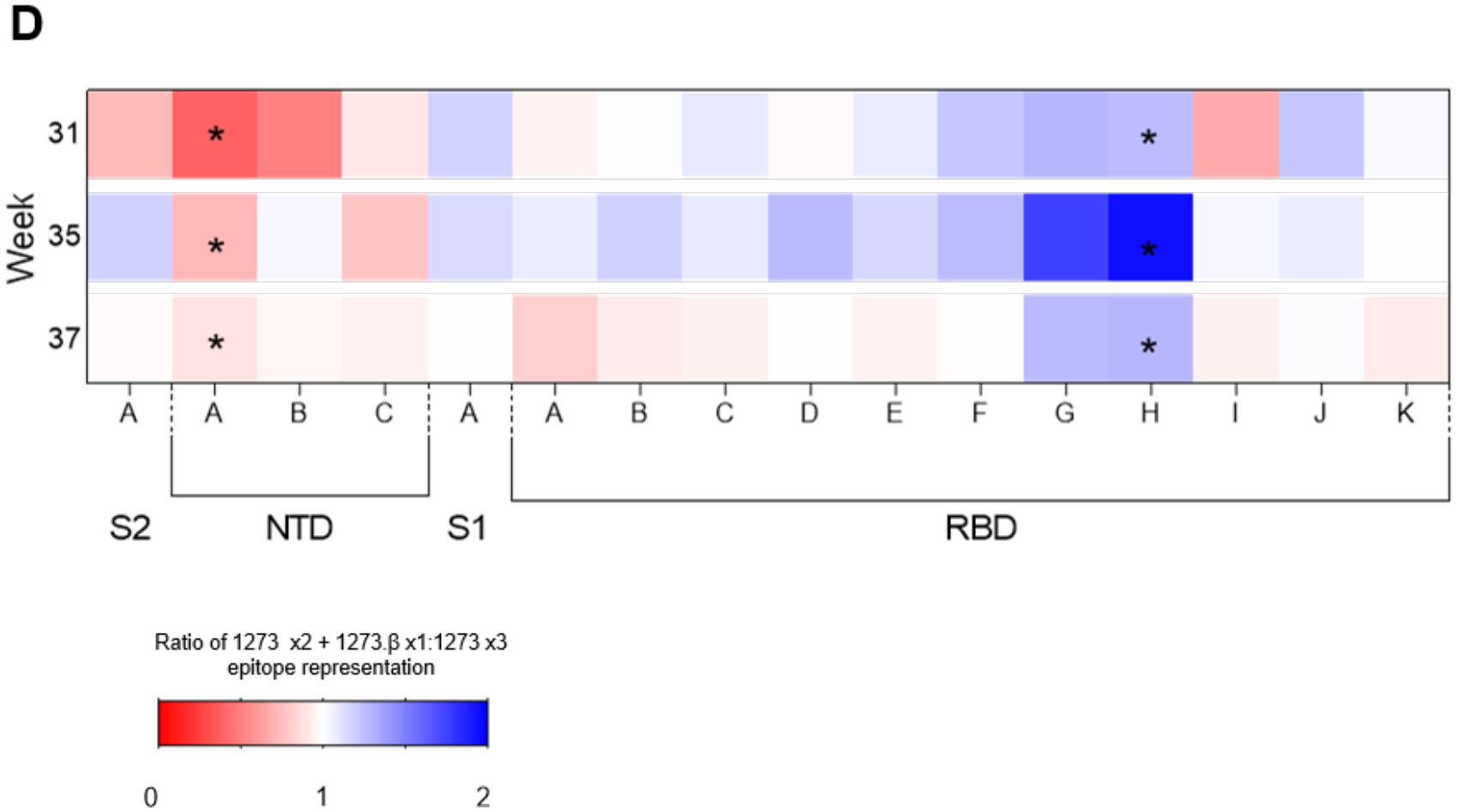
Serum Epitope Mapping Analysis. (A) Serum epitope fingerprints at Week 35 (6 weeks post-boost). Absolute serum epitope reactivity (RUs) is indicated from light to dark purple. Columns represent individual NHP, rows represent antigenic site. Bar graphs on the right depict relative serum reactivity (percent competition) for each antigenic site with each animal represented by an open circle (n=8/group). (B, C) Absolute (RUs) and relative (percent competition) serum reactivity for epitopes which show significant (p ≤ 0.05) differences between immunization groups at all time points. NTD-A site is represented by mAb 4-8, RBD site H is represented by mAb A23-97.1 *see Table S2*. (D) Ratio of mRNA-1273 x2 + mRNA-1273.β x1 to mRNA-1273 x3 epitope representation (percent competition) for each antigenic site at week 31, 35 and 37. Significant (p ≤ 0.05) differences determined by unpaired t-test between immunization groups at a single time point.

### Boosting expands mRNA-1273-induced memory B cell responses

The durability of vaccine-induced antibody responses is an integral component of the pandemic response as efforts continue to mitigate the ongoing spread of SARS-CoV-2. Durable humoral immunity is driven by the ability to generate and sustain memory B cells and long-lived plasmablasts (*28*), and recent data in humans shows the potential for such responses after vaccination with mRNA or primary infection (*29, 30*). To define how B cell specificity matures overtime in mRNA-1273 immunized NHP and to assess how mRNA-1273 priming imprints homologous or heterologous boost, we performed temporal analysis of WA-1 and β S-specific memory B cells (**Fig. S5**). Following the primary immunization series with mRNA-1273, the frequency of memory B cells expressing antibody receptors dual reactive for both WA-1 and β S at week 6 was 2-3%, with a much lower proportion of single WA-1 or β-specific B cells **(Fig. 4A-B)**. Six months later, at week 29, there was ∼10-fold reduction in the frequency of double-positive WA-1 and β S-specific memory B cells **(Fig. 4A-B)**, but these were restored (∼10-fold increase) following a memory boost with mRNA-1273 or mRNA-1273.β**(Fig. 4)**. In contrast, boosting did not cause an increase in the frequency of single-positive WA-1 or β S-specific B cells **(Fig. 4)**. This finding demonstrates a rapid recall response of primary vaccination B cells and coincides with an increase in neutralizing antibody responses **(Fig. 1E-J)**. The mRNA-1273.β immunized NHP had a memory B cell response that also consisted of WA-1 and β S-specific double-positive specificity, but a higher proportion of single positive β S-specific cells **(Fig. 4)**, consistent with β-specific skewing of antibody responses following mRNA-1273.β primary immunization **(Fig. S2)**. These data confirm the observation from serum antibody epitope mapping that both homologous and heterologous boosting can efficiently expand memory B-cell responses that are maintained after primary immunization.

**Figure 4.**
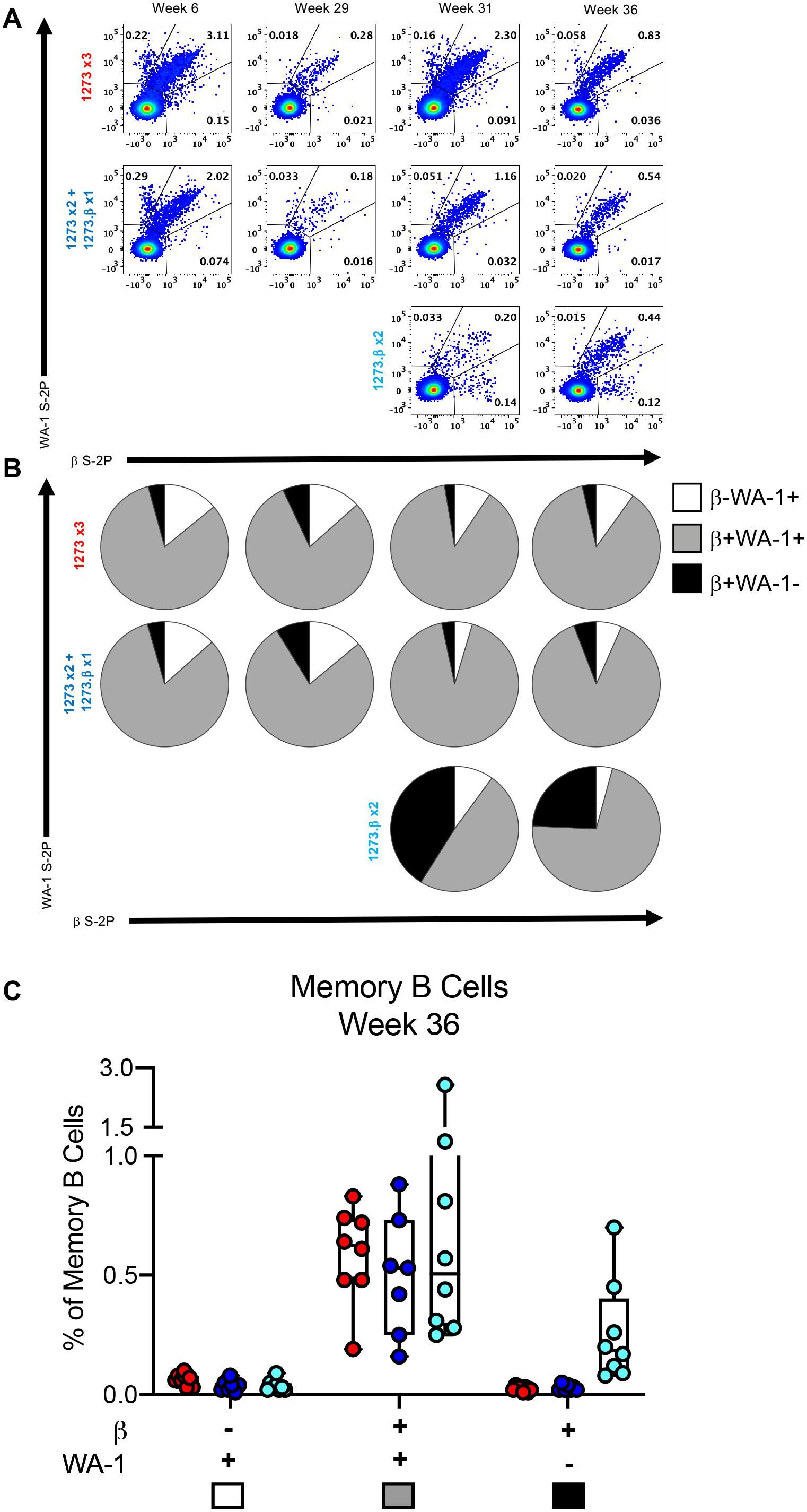
Frequency of SARS-CoV-2 spike-specific B cells in PBMC. (A) Representative flow cytometry plots from one NHP per vaccine group depicting the frequency of memory B cells that bind the WA-1 S, WA-1 S and β S, or β S at various time points after immunization. (B) Pie charts representing the fraction of S-specific B cells that are single positive for the WA-1 (white, β-WA-1+), double positive for WA-1 and β (gray, β+WA-1+), and single positive for β (black, b+WA-1-) for all NHP in each group. (C) Frequency of β-WA-1+, β+WA-1+, and β+WA-1-memory B cells at week 36 in each of the vaccine groups. Boxes and horizontal bars denote the IQR and medians, respectively; whisker end points are equal to the maximum and minimum values. Circles represent individual NHP.

### Boosting restimulates mRNA-1273-induced Th1 and Tfh T cell responses

mRNA-1273 has been shown to elicit CD4 responses comprised of Th1 and T follicular helper (Tfh) cells, and a lower frequency of CD8 T cells in both humans and NHP (*18, 19, 26, 31*). However, the definition of longitudinal T-cell development and the potential for boosting contracted memory T cells have not yet been reported in mRNA-1273 immunized NHP. Here, consistent with prior studies, S-specific Th1 and Tfh, but not Th2, responses were induced at week 6, 2 weeks following mRNA-1273 vaccination **(Fig. 5)**. In these NHP, Th1 and Tfh responses were contracted by week 29, but were both boosted by either mRNA-1273 or mRNA-1273.β **(Figs. 5A, and 5C-D)**. Week 36 responses revealed mRNA-1273.βimmunization also induced S-specific Th1 and Tfh, but not Th2, responses **(Fig. 5)**. These data suggest that homologous and heterologous mRNA boosts are equally capable of restimulating Th1 and Tfh T cell responses.

**Figure 5.**
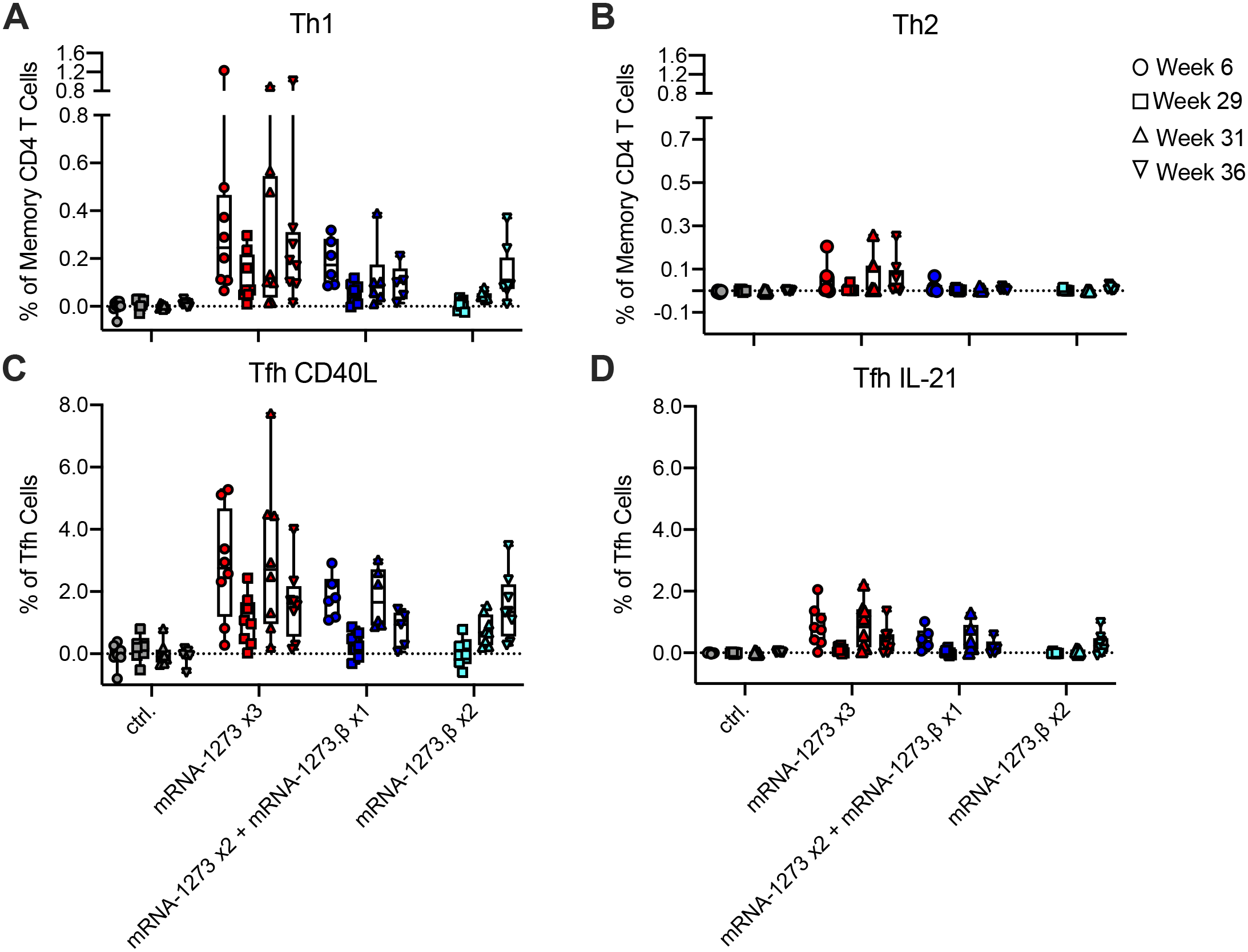
T cell responses following mRNA-1273 immunization. Rhesus macaques were immunized according to Figure S1. Intracellular staining (ICS) was performed on PBMCs at weeks 6, 29, 31, and 36 to assess T cell responses to SARS-CoV-2 S protein peptide pools, S1 and S2. Responses to S1 and S2 individual peptide pools were summed. (A) Th1 responses (IFNg, IL-2, or TNF), (B) Th2 responses (IL-4 or IL-13), (C) Tfh CD40L upregulation (peripheral follicular helper T cells (Tfh) were gated on central memory CXCR5^+^PD-1^+^ICOS^+^ CD4 T cells), (D) Tfh IL-21. Boxes and horizontal bars denote IQR and medians, respectively; whisker end points are equal to the maximum and minimum values. Circles represent individual NHP. Dotted lines are set to 0%.

### NHP are highly protected against SARS-CoV-2 β following a boost

NHP were challenged at week 38, ∼8 weeks post-boost in animals receiving three immunizations, and ∼5 weeks post primary mRNA-1273.β immunization (two immunizations), with a total dose of 2×10^5^ PFU of SARS-CoV-2 β by intratracheal (IT) and intranasal (IN) routes **(Fig. S1)**. Two days post-challenge, control NHP that received mock mRNA had a median of ∼6 log_10_ SARS-CoV-2 envelope (E) sgRNA (sgRNA _E) copies/mL in BAL. In contrast, all 8 NHP that received primary mRNA-1273.β immunization had undetectable BAL sgRNA_E. In NHP that received a boost after primary immunization with mRNA-1273, there was a hierarchy of protection against viral replication in the lung, where 4/8 NHP that received the homologous mRNA-1273 boost had undetectable BAL sgRNA _E, and 7/8 NHP boosted with heterologous mRNA-1273.β had undetectable BAL sgRNA_E. By day 4, all boosted animals with the exception of one mRNA-1273 boosted NHP had undetectable BAL sgRNA_E **(Fig. 6A)**. In NS, at days 2 and 4 post-challenge, both groups of boosted NHP showed significantly less (∼3-4 log_10_ reduction) sgRNA E compared to controls (*p* = 0.0014 and 0.0004 for mRNA-1273 and mRNA-1273.β, respectively). By day 7, the majority (13/16) of boosted NHP had undetectable sgRNA_E compared to control NHP, for which NS sgRNA_E persisted at a median of 4 log_10_ NS sgRNA_E (**Fig. 6B).**

**Figure 6.**
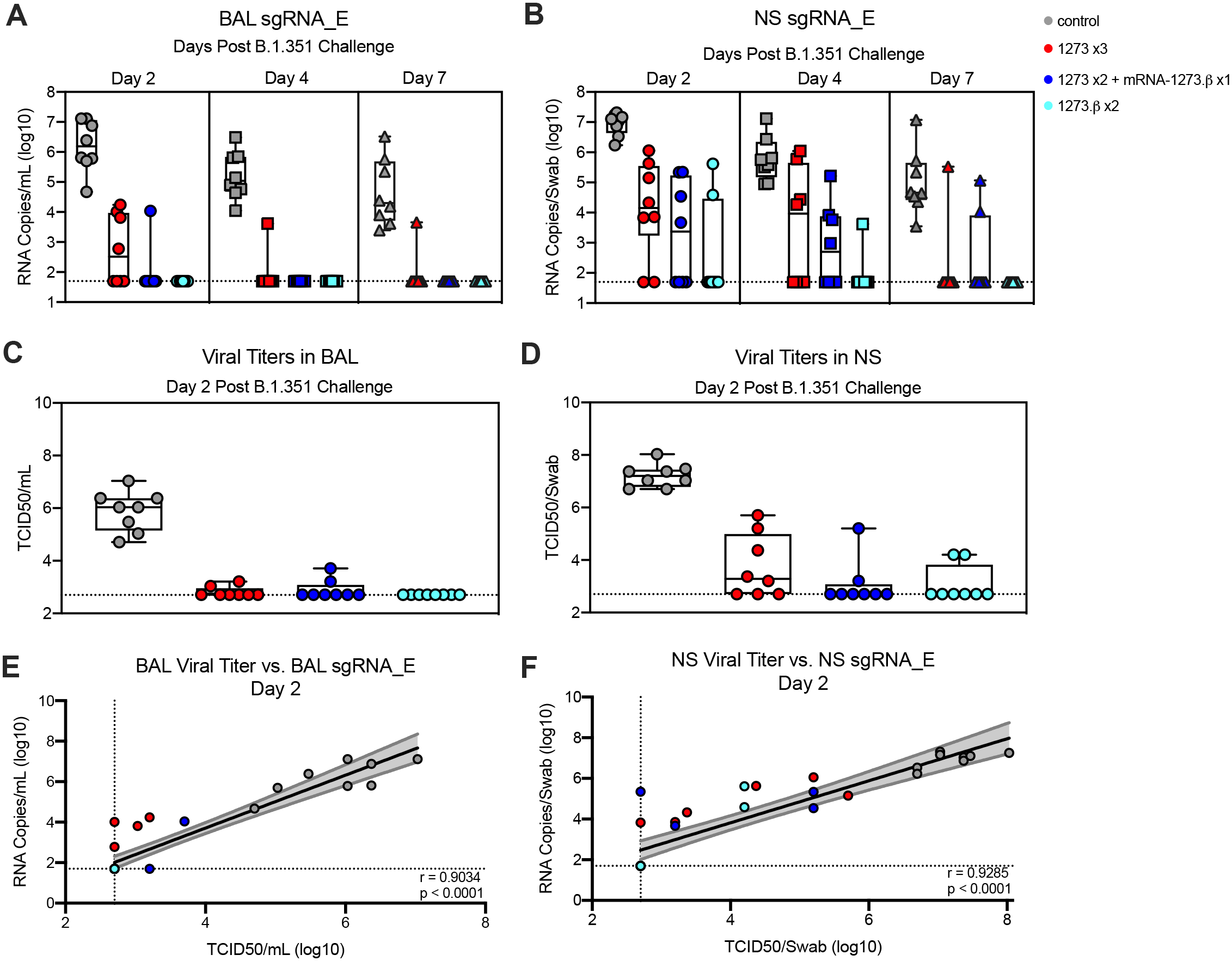
Efficacy of mRNA-1273 against upper and lower respiratory B.1.351 viral replication. Rhesus macaques were immunized and challenged as described in Figure S1. BAL (A, C) and nasal swabs (NS) (B, D) were collected on days 2 (circles), 4 (squares), and 7 (triangles) post-challenge, where applicable, and viral replication was assessed by detection of SARS-CoV-2_E sgRNA. Viral titers were assessed by TCID50 assay for BAL (C) and NS (D) collected on day 2 post-challenge. Boxes and horizontal bars denote the IQR and medians, respectively; whisker end points are equal to the maximum and minimum values. (E-F) Plots show correlations between viral titers and sgRNA_E in BAL (E) and NS (F) 2 days post-challenge. Black and gray lines indicate linear regression and 95% confidence interval, respectively. ‘r’ and ‘p’ represent Spearman’s correlation coefficients and corresponding p-values, respectively. Symbols represent individual NHP and may overlap for equal values. Dotted lines indicate assay limits of detection.

While sgRNA is a sensitive measurement of viral replication that can be used diagnostically, we also determined the ability of live virus to be propagated from post-challenge BAL and NS samples, an alternative measure of viral load that is relevant for indicating lung disease or transmission from the upper airway. On day 2 post-challenge, all boosted animals had low to undetectable (<4 log_10_ TCID_50_/mL) of culturable SARS-CoV-2 in BAL **(Fig. 6C)**. In NS, SARS-CoV-2 was unculturable for 3/8 and 5/8 NHP that were boosted with mRNA-1273 or mRNA-1273.β, respectively, which was similar to the mRNA-1273.β primary immunization group **(Fig. 6D)**. sgRNA and viral titers were highly correlated **(Fig. 6E-F)**; in fact, there was no culturable SARS-CoV-2 from BAL or NS that had sgRNA_E levels <1.0 x10^4^ RNA copies/mL **(Fig. 6E)** and <4.7 x 10^3^ RNA copies/swab **(Fig. 6F)**, respectively. Nucleocapsid (N)-specific sgRNA measurements followed the same trend, albeit to with higher detection sensitivity **(Fig. S7)**.

We also evaluated lung samples for pathology and detection of viral antigen (VAg) 7-9 days post SARS-CoV-2 β challenge. Inflammation was minimal to mild and was similar across lung samples from vaccinated NHP, with rare cases showing a moderate to severe response **(Fig. S8 and Table S3)**. The inflammatory changes in the lung were characterized by a mixture of macrophages and polymorphonuclear cells present within some alveolar spaces and mild to moderate expansion of alveolar capillaries with mild Type II pneumocyte hyperplasia at day 7 post-challenge to changes more consistent with lymphocytes, histiocytes and fewer polymorphonuclear cells associated with more prominent and expanded alveolar capillaries, occasional areas of perivascular and peribronchiolar inflammation, and Type II pneumocyte hyperplasia at days 8-9 post-challenge. SARS-CoV-2 antigen (Ag) was detected in 6/6 control NHP evaluated at days 7, 8, or 9 after infectious challenge, with 3/6 showing Ag present in multiple lobes. In vaccinated NHP, Ag was detected in only 1/24 animals, and in a single lobe (mRNA-1273×3, **Table S3)**. These results, together with sgRNA data, confirm that boosting mRNA-1273 immunized NHP limits βviral replication in the lower and upper airway.

## DISCUSSION

The SARS-CoV-2 mRNA-1273 vaccine shows between 90-100% protection against WA-1 (*2*), α, or β variants (*32*) when administered as two doses four weeks apart and assessed within a 2 month window. Kinetic analyses of antibody responses following vaccination with mRNA-1273 (*7*) or BNT162b2 (*30*) show reduction of neutralizing activity from the peak humoral response after the second immunization through day 209 (*7, 17*). Consistently, the β variant is the most neutralization resistant of all VOC to date. However, it has not yet been established how this resistance affects the durability of protective efficacy in humans. Recent clinical reports from Israel show that cohorts immunized with the BioNTech/Pfizer vaccine over 6 months ago may present with symptomatic disease when infected with VOC (*33*). These data suggest that it may be important to boost antibody responses, especially against VOC, to sustain protection against severe disease, especially in at-risk cohorts and to reduce the potential for transmission. A related issue is whether the additional boost should be matched to a specific newly arising variant (heterologous) or if homologous boosting with the original vaccine based on ancestral strains will be sufficient to generate broadly neutralizing antibody responses against VOC. Here, we show that boosting NHP ∼ 6 months after their primary vaccination series with either mRNA-1273 or mRNA-1273.β significantly increased serum neutralizing activity against all variants and mediated high-level protection in the upper and lower airways against challenge with the β variant.

Longitudinal neutralizing antibody responses against the benchmark D614G strain compared to β and several additional VOC confirmed data from prior vaccine studies in humans or NHP (*7, 30, 34*), that there was a 3-10 fold reduction in neutralizing activity against several VOC, with β and γ variants showing the greatest reduction at the peak time point after the primary immunization series. Here, all neutralizing responses were significantly reduced by week 24. Of note, there was a more rapid decay of binding antibodies to VOCs compared to WA-1, but a relatively slower decay in serum neutralizing activity against VOC. These data suggest that, although binding antibody is an established correlate for short term protection, deeper analysis is needed to define correlates of protection with contracted memory responses. These data also raise a possibility that while there is a quantitative decrease in the magnitude of antibody responses over time following vaccination, the loss of functional activity may be partially mitigated by affinity maturation and improved quality of the response. It was recently reported that there may be discordance in antibody evolution between convalescent individuals and those who receive an mRNA vaccine. Specifically, less affinity maturation and a limited increase in breadth to the RBD subdomain was observed in isolated monoclonal antibodies between 2- and 5-months post mRNA vaccination (*30*). Our assessment of polyclonal serum antibody affinities (avidity) using the intact WA-1 S protein showed that vaccination with mRNA-1273 led to an increase in avidity over time, confirming a qualitative improvement in the antibody response in NHP (*29, 35, 36*).

Following a homologous mRNA-1273, or heterologous mRNA-1273.β boost at 6 months after the primary immunization series, there was a ∼7-21-fold increase in serum antibody neutralizing activity for D614G, β, and the other VOCs tested, and the increase was not significantly different between homologous or heterologous boosting. These data are consistent with recent studies in humans showing that boosting with mRNA-1273 or mRNA-1273.β approximately 6 months after the primary immunization with mRNA-1273 elicited significantly higher neutralizing titers against the D614G, β, and the other VOC tested compared to pre-boost titers (*37*). Importantly, there was significant boosting of neutralizing responses against the β variant even in individuals that had undetectable responses prior to the boost. Last, neutralizing responses after the boost in our NHP study and in humans were significantly higher than the peak titers observed after the primary vaccination series (*17*).

The increase in antibody responses post-boost suggests that there is significant B-cell memory induced by primary mRNA vaccination that can be rapidly recalled following the boost (*7, 29*). Here, we show that after primary immunization with mRNA-1273, the peak response yields a total frequency of WA-1 and β-specific memory B cells of ∼ 3%, and the majority (∼ 80%) of these responses recognized both S proteins, with a small proportion specific for WA-1 only. There was a very low frequency of B cells specific for β only. After 6 months, all responses contracted to exhibit a 10-fold decrease in the frequency of WA-1 and β-specific memory B cells, but these were rapidly restored to a frequency of 3% after boosting. The relative frequency of B cells specific for WA-1, β, or both did not change post-boost suggesting that priming with mRNA-1273 imprinted the B-cell repertoire.

The boosting of B cells specific for WA-1 and β highlights the overlapping epitopes in S between the two strains and likely accounts for neutralization breadth against VOCs. A notable finding was a limited (<5%) frequency of B cells specific for β only following mRNA-1273 vaccination or after the homologous or heterologous boost. However, naïve animals primed with only mRNA-1273.β showed between 25-40% of the total B-cell response specific for β only. Likewise, serum antibody epitope profiling demonstrated a repertoire that was qualitatively similar following either homologous or heterologous memory boost after mRNA-1273 primary vaccination, while a primary vaccination series with mRNA-1273.β yielded a unique repertoire (both qualitative and quantitative) with marked increases in reactivity to epitopes associated with broad and potent neutralization of VOCs. These data have implications for how future mRNA vaccines can be designed to imprint B-cell repertoires in naïve individuals for increased potency and breadth of neutralizing activity (*38*).

While the data presented here focus on the role of boosting vaccine responses with mRNA, as a proof-of-principle we also show that priming with mRNA-1273.β in naïve NHP induced potent neutralizing antibody responses against β and provided high-level protection in the upper and lower airways following challenge. Parallel studies in pre-clinical rodent models have shown that priming with a variant mRNA or a bi-valent vaccine containing the WA-1 and the β variant induces higher neutralizing responses to D614G and a number of variants tested relative to either mRNA-1273 or mRNA-1273.β alone (*38*).

In conclusion, the data reported here show that a booster dose can significantly improve the breadth and potency of neutralizing antibody responses sufficient to mediate protection against upper and lower airway infection by a heterologous challenge virus shown to be relatively resistant to *in vitro* neutralization. The potential clinical utility of a boost would be to sustain high-level protection against severe disease and possibly limit the duration and extent of mild infection in the setting of waning immunity especially in the elderly, and other groups with pre-existing health conditions or poor response to vaccination. We have previously shown that there is a higher threshold for antibody-mediated protection in the upper airway (*18*), which would be more important for limiting mild infection and transmission than for protecting the lower airways to limit severe disease. We speculate that new variants with high viral load and transmissibility, such as δ, may require a late boost to increase antibody responses in the upper airway enough to limit transmission. Furthermore, if such variants have more rapid progression to lower airway disease than prior strains, boosting may be more important for mitigating severe disease as serum antibody wanes, because disease progression may outpace otherwise protective anamnestic responses. Last, a late boost will potentially provide more durable and broader immunity and would reduce morbidity and mortality until greater population immunity is achieved.

## MATERIALS AND METHODS

### Pre-clinical mRNA and lipid nanoparticle production

A sequence-optimized mRNA encoding prefusion-stabilized SARS-CoV-2 S-2P protein (*39, 40*) for WA-1 and β was synthesized *in vitro* and formulated as previously reported (*17, 41*). Control mRNA “UNFIX-01 (Untranslated Factor 9)” was synthesized based on the sequence in **Table S4** and similarly formulated into lipid nanoparticles.

### Rhesus macaque model

Animal experiments were carried out in compliance with all pertinent US National Institutes of Health regulations and approval from the Animal Care and Use Committees of the Vaccine Research Center and Bioqual, Inc. (Rockville, MD). Studies were conducted at Bioqual, Inc. The experimental details of VRC-20-857.3b **(Fig. S1)** are similar to prior studies (*18, 19, 26, 34*). Briefly, 3–15-year-old rhesus macaques of Indian origin were stratified into groups based sex, age, and weight. Animals were immunized with mRNA-1273 at week 0 and week 4 with a dose of 100 µg intramuscularly (IM) in 1 ml formulated in PBS into the right hindleg. Placebo-control animals were administered control mRNA. At week 29 (∼ 25 weeks after the second immunization), a group of animals were boosted with 50 µg of mRNA-1273 or 50 µg of mRNA-1273.β. An additional group of animals were immunized at week 29 and 33 with 50 µg of mRNA-1273.β. At week 38 (9 weeks after the homogolous or heterologous mRNA boost or 5 weeks after the mRNA-1273.β prime and boost) all animals were challenged with a total dose of 2×10^5^ PFU SARS-CoV-2 β (JHU P2) as previously described (*34*). The viral inoculum was administered as 1.5×10^5^ PFU in 3 mL intratracheally (IT) and 0.5×10^5^ PFU in 1 mL intranasally (IN) in a volume of 0.5 mL into each nostril. Pre- and post-challenge sample collection is detailed in **Fig. S1**.

### Quantification of SARS-CoV-2 subgenomic RNA (sgRNA)

BAL and nasal swab (NS) subgenomic SARS-CoV-2 E mRNA was quantified via reverse transcription-polymerase chain reaction (RT-PCR) as previously described (*26*). Subgenomic SARS-CoV-2 N mRNA was quantified similarly, as described in (*34*). The lower limit of quantification was 50 copies.

### TCID50 quantification of SARS-CoV-2 from BAL

Viral load (TCID50) from BAL samples was calculated using previously described methods (*34*).

### Histopathology and immunohistochemistry (IHC)

Post-challenge, on days 7-9, animals were euthanized, and lung tissue was processed and stained with hematoxylin and eosin (H&E) for routine histopathology and analyzed for detection of SARS-CoV-2 virus antigen as previously described (*34*). All samples were blinded and evaluated by a board-certified veterinary pathologist.

### Multiplex meso-scale ELISA for serum antibody responses

For 10-plex ELISA, 96-well plates precoated with SARS-CoV-2 S-2P (*39*) and RBD proteins from multiple variants, SARS-CoV-2 N protein, and Bovine Serum Albumin (BSA) and supplied by the manufacturer [Meso Scale Display (MSD)]. Determination of serum antibody binding was performed as previously described (*7*), and reagent details are in **Table S5**. All calculations are performed within Excel and the GraphPad Prism software, v7.0. Readouts are provided as Area Under Curve (AUC). For 4-plex ELISA, 96-well plates were pre-coated with SARS-CoV-2 S-2P, RBB, and N and a BSA control in each well. Determination of serum antibody binding was performed as previously described (*7, 18, 26*). Calculated ECLIA parameters to measure binding antibody activities included interpolated concentrations or assigned arbitrary units (AU/ml) read from the standard curve. International units (IU) were established for each antigen based on parallelism between MSD reference standard and the WHO international standard. S-specific IgG lower limit of detection = 0.3076 IU/mL. RBD-specific IgG lower limit of detection = 1.5936 IU/mL.

### Meso-scale ELISA for mucosal antibody responses

Total S-specific IgG in BAL and nasal washes was determined by meso-scale ELISA (MSD) as previously described (*46*).

### Serum antibody avidity assay

Avidity was assessed using a sodium thiocyanate (NaSCN)-based avidity ELISA against SARS-CoV-2 S-2P as previously described (*18, 26*). The avidity index (AI) was calculated using the ratio of IgG binding to S-2P in the absence or presence of NaSCN and reported as the average of two independent experiments, each containing duplicate samples.

### Serum antibody epitope definition

Serum epitope mapping competition assays were performed using a Biacore 8K+ (Cytiva) surface plasmon resonance (SPR) spectrometer. According to manufacturer’s protocol, anti-histidine IgG1 antibody was immobilized on Series S Sensor Chip CM5 (Cytiva) through primary amine coupling using a His capture kit (Cytiva). His-tagged SARS-CoV-2 WA.1 S protein containing 2 proline stabilization mutations (S-2P) was captured on active sensor surface.

Competitor monoclonal antibody (mAb) or a negative control antibody at set concentrations were injected over both active and reference surfaces to saturation. Competitor human IgG mAbs used include: S2-specific mAb S652-112, NTD-specific mAbs 4-8, S652-118, and N3C, S1-specific mAb A20-36.1 and RBD-specific mAbs B1-182, CB6, A20-29.1, A19-46.1, LY-COV555, A19-61.1, S309, A23-97.1, A19-30.1, A23-80.1 and CR3022. Non-human primate sera were flowed over both active and reference sensor surfaces for 40 minutes. 1X PBS-P+ (Cytiva) was used as the running buffer and diluent for all samples. Active and reference sensor surfaces were regenerated between each analysis cycle using 10mM glycine, pH 1.5 (Cytiva).

Prior to analysis, sensorgrams were aligned to Y (Response Units) = 0, beginning at the serum association phase using Biacore 8K Insights Evaluation Software (Cytiva). Reference-subtracted relative “analyte binding late” report points (RU) were collected and used to calculate absolute and relative competition. Relative percent competition (% C) was calculated using the following formula: % C = [1 – (100 * ( (RU in presence of competitor mAb) / (RU in presence of negative control mAb))]. Absolute competition (ΔRUs) was calculated with the following formula: ΔRUs = [(RU in presence of negative control mAb) – (RU in presence of competitor mAb)]. Results are reported as absolute serum epitope reactivity and percent competition and statistical analysis was performed using unpaired, two-tailed t-test (Graphpad Prism v.8). All assays were performed in duplicate, with average data point for each animal represented on corresponding graphs.

### Surface plasmon resonance binding assay

Series S CM5 Sensor Chips (Cytiva) were activated by immobilizing anti-histidine IgG1 antibody on surface using a His capture kit (Cytiva), according to manufacturer’s protocol. His-tagged SARS-CoV-2 WA.1 or β S protein containing 2 proline stabilization mutations (S-2P) was captured on active sensor surface at a set concentration for 10 minutes. Monoclonal antibodies (mAbs) were injected over both active and reference surfaces and allowed to bind to saturation. Graphs represent reference-subtracted relative “analyte binding late” report points (RUs), determined by aligning sensorgrams to Y (Response Units) = 0, beginning at the mAb association stage using Biacore 8K Insights Evaluation Software (Cytiva). All assays were performed using a Biacore 8K+ (Cytiva) surface plasmon resonance (SPR) spectrometer.

### ACE2-binding inhibition assay

ACE2-binding inhibition was completed, as previously described (*47*), using 1:40 diluted BAL and nasal wash samples.

### Lentiviral pseudovirus neutralization assay

Pseudotyped lentiviral reporter viruses were produced by the co-transfection of plasmids encoding SARS-CoV-2 proteins from multiple variants, a luciferase reporter, lentivirus backbone, and human transmembrane protease serine 2 (TMPRSS2) genes as previously described (*17, 37*); reagent details are in **Table S5**. Sera were tested, in duplicate, for neutralizing activity against the pseudoviruses and percent neutralization was calculated in Prism v9.0.2 (GraphPad). The lower limit of quantification was 1:40 ID_50_.

### VSV pseudovirus neutralization assay

SARS-CoV-2 pseudotyped recombinant VSV-ΔG-firefly luciferase viruses were made by co-transfection of plasmid expressing full-length S, and subsequent infection with VSVΔG-firefly-luciferase and neutralization assays were completed on sera samples as previously described (*15, 42*); reagent details are in **Table S5**. The lower limit of quantification was 1:40 ID_50_.

### Focus reduction neutralization test (FRNT)

FRNT assays were performed on sera samples, in duplicate, as previously described (*34*); and reagent details are in **Table S5**. For samples that do not neutralize 50% of virus at the limit of detection, 5 was plotted and used for geometric mean calculations.

### B cell Probe Binding Assay

Cryopreserved PBMC were thawed and washed in wash buffer (4% Heat-inactivated newborn calf serum/0.02% NaN_3_/ phenol-free RPMI). Cells were incubated with streptavidin-BV605 (BD Biosciences) labeled β S-2P and streptavidin-BUV661 (BD Biosciences) labeled WA-1 S-2P for 30 minutes at 4°C, washed twice in 1X PBS, and incubated with Aqua live/dead fixable dead cell stain (Thermo Fisher Scientific) for 20 minutes at room temperature. After two washes in wash buffer, cells were incubated with primary antibodies for 20 minutes at room temperature. The following antibodies were used (monoclonal unless indicated): IgD FITC (goat polyclonal, Southern Biotech), IgM PerCP-Cy5.5 (clone G20-127, BD Biosciences), IgA Dylight 405 (goat polyclonal, Jackson Immunoresearch Inc), CD20 BV570 (clone 2H7, Biolegend), CD27 BV650 (clone O323, Biolegend), CD14 BV785 (clone M5E2, Biolegend), CD8 BUV395 (clone RPA-T8, BD Biosciences), CD16 BUV496 (clone 3G8, BD Biosciences), CD4 BUV737 (clone SK3, BD Biosciences), CD19 APC (clone J3-119, Beckman Coulter), IgG Alexa 700 (clone G18-145, BD Biosciences), CD3 APC-Cy7 (clone SP34.2, BD Biosciences), CD38 PE (clone OKT10, Caprico Biotechnologies), CD21 PE-Cy5 (clone B-ly4, BD Biosciences), and CXCR5 PE-Cy7 (clone MU5UBEE, Thermo Fisher Scientific). Cells were washed twice in wash buffer and residual red blood cells were lysed using BD FACS Lysing Solution (BD Biosciences) for 10 minutes at room temperature. Following two additional washes, cells were fixed in 0.5% formaldehyde (Tousimis Research Corp.) All antibodies were previously titrated to determine the optimal concentration. Samples were acquired on an BD FACSymphony flow cytometer and analyzed using FlowJo version 10.7.2 (BD, Ashland, OR).

### Intracellular cytokine staining

Cryopreserved PBMC were thawed and rested overnight in a 37C/5% CO2 incubator. The next morning, cells were stimulated with SARS-CoV-2 Spike protein peptide pools (S1 and S2) that were matched to the vaccine insert (comprised of 158 and 157 individual peptides, respectively, as 15mers overlapping by 11 aa in 100% DMSO, JPT Peptides) at a final concentration of 2 μg/ml in the presence of 3 mM monensin for 6 hours. Negative controls received an equal concentration of DMSO instead of peptides (final concentration of 0.5%). Intracellular cytokine staining was performed as described (*43*). The following monoclonal antibodies were used: CD3 APC-Cy7 (clone SP34.2, BD Biosciences), CD4 PE-Cy5.5 (clone S3.5, Invitrogen), CD8 BV570 (clone RPA-T8, Biolegend), CD45RA PE-Cy5 (clone 5H9, BD Biosciences), CCR7 BV650 (clone G043H7, Biolegend), CXCR5 PE (clone MU5UBEE, Thermo Fisher), CXCR3 BV711 (clone 1C6/CXCR3, BD Biosciences), PD-1 BUV737 (clone EH12.1, BD Biosciences), ICOS Pe-Cy7 (clone C398.4A, Biolegend), CD69 ECD (cloneTP1.55.3, Beckman Coulter), IFN-g Ax700 (clone B27, Biolegend), IL-2 BV750 (clone MQ1-17H12, BD Biosciences), IL-4 BB700 (clone MP4-25D2, BD Biosciences), TNF-FITC (clone Mab11, BD Biosciences), IL-13 BV421 (clone JES10-5A2, BD Biosciences), IL-17 BV605 (clone BL168, Biolegend), IL-21 Ax647 (clone 3A3-N2.1, BD Biosciences), and CD154 BV785 (clone 24-31, BioLegend). Aqua live/dead fixable dead cell stain kit (Thermo Fisher Scientific) was used to exclude dead cells. All antibodies were previously titrated to determine the optimal concentration. Samples were acquired on an BD FACSymphony flow cytometer and analyzed using FlowJo version 9.9.6 (Treestar, Inc., Ashland, OR). Gating schema is represented in **Fig. S6**.

### Statistical Analysis

Graphs show data from individual animals with dotted lines indicating assay limits of detection. Groups were compared for viral load and immune responses using Welch’s t-tests; data were analyzed on the log10 scale for viral loads and appropriate immune assays. Changes over time were summarized using the change on the log_10_ scale, and statistical significance was determined using paired t-tests. When more than two variants are compared, as in Figure 1 A-F, a Holm’s adjustment for multiple comparisons was used on the set of p-values to determine significance; there were no adjustments for multiple comparisons across different analyses in this descriptive study. Comparisons drawn between vaccination groups for serum antibody epitope analysis were performed using unpaired *t* tests, within each time point evaluated. Analyses were performed in R version 4.0.2 and Prism (GraphPad) versions 8.2 and 9.0.2.

## Acknowledgements

We thank J. Stein and M. Young for technology transfer and administrative support, respectively. We thank members of the NIH NIAID VRC Translational Research Program, including H. Bao, E. McCarthy, J. Noor, A. Taylor, and R. Woodward, for technical and administrative assistance with animal experiments. We thank H. Mu and M. Farzan for the ACE2-overexpressing 293 cells. We thank B. Zhang for production and purification of published mAbs. We thank T. Ruckwardt, E. Phung, I.-T. Teng, A. Olia, O. Abiona and A. DiPiazza for contributions to probe production and validation. We thank M. Brunner and M. Whitt for kind support on recombinant VSV-based SARS-CoV-2 pseudovirus production.

## Author Contributions

K.S.C., M.G., D.W., S.O., S.R.N., D.R.F., S.F.A., R.L.D., B.F., T.S.J., C.S., L.L., D.V., A.V.R., Z.F., A.P.W., J.I.M., M.S., S.O., S.D.S, C.T., A.C., M.K., K.W.B., M.M., B.M.N., G.S.A., A.R.H., F.L., C.S., C.S., L.W., E.L., S.T.N., S.J.P., M.M.D., J.M., J-P.M.T., A.C., A.D., N.D., A.P., L.P., H.A., K.E.F., J.M., K.W., D.K.E., M.C.N., J.M., I.N.M., A.C., M.G.L., M.S.S., M.R., A.M., D.C.D., N.J.S., B.S.G., and R.A.S. designed, completed, analyzed, and/or supervised experiments. Y.Z., L.W., S.B-B., M.C., W.S., A.P., E.B., provided critical published reagents/analytic tools. K.S.C., M.C.N., N.J.S., M.R., B.S.G, and R.A.S. wrote the manuscript. K.S.C., D.W., M.S., K.E.F., S.F.A., and G.A. prepared figures and tables. All authors contributed to discussions about and editing of the manuscript.

## Data and materials availability

All data are available in the main text or the supplementary materials.

## References

1. F. P. Polack et al., Safety and Efficacy of the BNT162b2 mRNA Covid-19 Vaccine. New England Journal of Medicine 383, 2603–2615 (2020).

2. L. R. Baden et al., Efficacy and Safety of the mRNA-1273 SARS-CoV-2 Vaccine. New England Journal of Medicine 384, 403–416 (2020).

3. M. Voysey et al., Safety and efficacy of the ChAdOx1 nCoV-19 vaccine (AZD1222) against SARS-CoV-2: an interim analysis of four randomised controlled trials in Brazil, South Africa, and the UK. The Lancet 397, 99–111 (2021).

4. J. Sadoff et al., Safety and Efficacy of Single-Dose Ad26.COV2.S Vaccine against Covid-19. New England Journal of Medicine, (2021).

5. P. T. Heath et al., Safety and Efficacy of NVX-CoV2373 Covid-19 Vaccine. New England Journal of Medicine, (2021).

6. C. Liu et al., Reduced neutralization of SARS-CoV-2 B.1.617 by vaccine and convalescent serum. Cell, (2021).

7. A. Pegu et al., Durability of mRNA-1273-induced antibodies against SARS-CoV-2 variants. bioRxiv, 2021.2005.2013.444010 (2021).

8. Q. Li et al., The Impact of Mutations in SARS-CoV-2 Spike on Viral Infectivity and Antigenicity. Cell 182, 1284–1294 e1289 (2020).

9. H. Tegally et al., Detection of a SARS-CoV-2 variant of concern in South Africa. Nature, (2021).

10. C. K. Wibmer et al., SARS-CoV-2 501Y.V2 escapes neutralization by South African COVID-19 donor plasma. Nat Med, (2021).

11. E. Andreano et al., SARS-CoV-2 escape in vitro from a highly neutralizing COVID-19 convalescent plasma. bioRxiv, (2020).

12. M. Hoffmann et al., SARS-CoV-2 variants B.1.351 and P.1 escape from neutralizing antibodies. Cell, (2021).

13. W. F. Garcia-Beltran et al., Multiple SARS-CoV-2 variants escape neutralization by vaccine-induced humoral immunity. Cell, (2021).

14. N. Doria-Rose et al., Antibody Persistence through 6 Months after the Second Dose of mRNA-1273 Vaccine for Covid-19. N Engl J Med, (2021).

15. K. Wu et al., Serum Neutralizing Activity Elicited by mRNA-1273 Vaccine. New England Journal of Medicine 384, 1468–1470 (2021).

16. T. Tada et al., Comparison of Neutralizing Antibody Titers Elicited by mRNA and Adenoviral Vector Vaccine against SARS-CoV-2 Variants. bioRxiv, 2021.2007.2019.452771 (2021).

17. K. Wu et al., Preliminary Analysis of Safety and Immunogenicity of a SARS-CoV-2 Variant Vaccine Booster. medRxiv, 2021.2005.2005.21256716 (2021).

18. K. S. Corbett et al., Immune correlates of protection by mRNA-1273 vaccine against SARS-CoV-2 in nonhuman primates. Science, eabj0299 (2021).

19. K. S. Corbett et al., Evaluation of the mRNA-1273 Vaccine against SARS-CoV-2 in Nonhuman Primates. New England Journal of Medicine 383, 1544–1555 (2020).

20. M. Guebre-Xabier et al., NVX-CoV2373 vaccine protects cynomolgus macaque upper and lower airways against SARS-CoV-2 challenge. Vaccine 38, 7892–7896 (2020).

21. N. B. Mercado et al., Single-shot Ad26 vaccine protects against SARS-CoV-2 in rhesus macaques. Nature 586, 583–588 (2020).

22. J. Yu et al., DNA vaccine protection against SARS-CoV-2 in rhesus macaques. Science 369, 806 (2020).

23. P. J. Klasse, D. F. Nixon, J. P. Moore, Immunogenicity of clinically relevant SARS-CoV-2 vaccines in nonhuman primates and humans. Science Advances 7, eabe8065 (2021).

24. N. van Doremalen et al., ChAdOx1 nCoV-19 vaccine prevents SARS-CoV-2 pneumonia in rhesus macaques. Nature 586, 578–582 (2020).

25. E. S. Winkler et al., Human neutralizing antibodies against SARS-CoV-2 require intact Fc effector functions for optimal therapeutic protection. Cell 184, 1804–1820.e1816 (2021).

26. J. R. Francica et al., Protective antibodies elicited by SARS-CoV-2 spike protein vaccination are boosted in the lung after challenge in nonhuman primates. Science Translational Medicine, eabi4547 (2021).

27. L. Wang et al., Ultrapotent antibodies against diverse and highly transmissible SARS-CoV-2 variants. Science, (2021).

28. G. D. Victora, M. C. Nussenzweig, Germinal Centers. Annual Review of Immunology 30, 429–457 (2012).

29. J. S. Turner et al., SARS-CoV-2 infection induces long-lived bone marrow plasma cells in humans. Nature 595, 421–425 (2021).

30. A. Cho et al., Antibody Evolution after SARS-CoV-2 mRNA Vaccination. bioRxiv, 2021.2007.2029.454333 (2021).

31. N. Pardi et al., Nucleoside-modified mRNA vaccines induce potent T follicular helper and germinal center B cell responses. The Journal of experimental medicine 215, 1571–1588 (2018).

32. H. Chemaitelly et al., mRNA-1273 COVID-19 vaccine effectiveness against the B.1.1.7 and B.1.351 variants and severe COVID-19 disease in Qatar. Nat Med, (2021).

33. S. Gazit et al., BNT162b2 mRNA Vaccine Effectiveness Given Confirmed Exposure; Analysis of Household Members of COVID-19 Patients. medRxiv, 2021.2006.2029.21259579 (2021).

34. K. S. Corbett et al., Evaluation of mRNA-1273 against SARS-CoV-2 B.1.351 Infection in Nonhuman Primates. bioRxiv, 2021.2005.2021.445189 (2021).

35. C. Gaebler et al., Evolution of antibody immunity to SARS-CoV-2. Nature 591, 639–644 (2021).

36. Z. Wang et al., Naturally enhanced neutralizing breadth against SARS-CoV-2 one year after infection. Nature 595, 426–431 (2021).

37. L. A. Jackson et al., An mRNA Vaccine against SARS-CoV-2 - Preliminary Report. N Engl J Med 383, 1920–1931 (2020).

38. C. A. Wu K, Koch M, Elbashir MS, Ma LZ, Lee D, Woods A, Henry C, Palandjian C, Hill A, Quinones J, Nunna N, O’Connell S, McDermott AB, Falcone S, Narayanan E, Colpitts T, Variant SARS-CoV-2 mRNA vaccines confer broad neutralization as primary or booster series in mice. bioRxiv, (2021).

39. D. Wrapp et al., Cryo-EM structure of the 2019-nCoV spike in the prefusion conformation. Science 367, 1260–1263 (2020).

40. J. Pallesen et al., Immunogenicity and structures of a rationally designed prefusion MERS-CoV spike antigen. Proceedings of the National Academy of Sciences 114, E7348–E7357 (2017).

41. K. J. Hassett et al., Optimization of Lipid Nanoparticles for Intramuscular Administration of mRNA Vaccines. Molecular Therapy - Nucleic Acids 15, 1–11 (2019).

42. M. A. Whitt, Generation of VSV pseudotypes using recombinant ΔG-VSV for studies on virus entry, identification of entry inhibitors, and immune responses to vaccines. J Virol Methods 169, 365–374 (2010).

43. M. M. Donaldson, S. F. Kao, K. E. Foulds, OMIP-052: An 18-Color Panel for Measuring Th1, Th2, Th17, and Tfh Responses in Rhesus Macaques. Cytometry A 95, 261–263 (2019).

