## Supplemental Material for "Protection against SARS-CoV-2 Beta Variant in mRNA-1273 Boosted Nonhuman Primates"

**This PDF file includes:**

Figs. S1 to S8
Tables S1 to S5

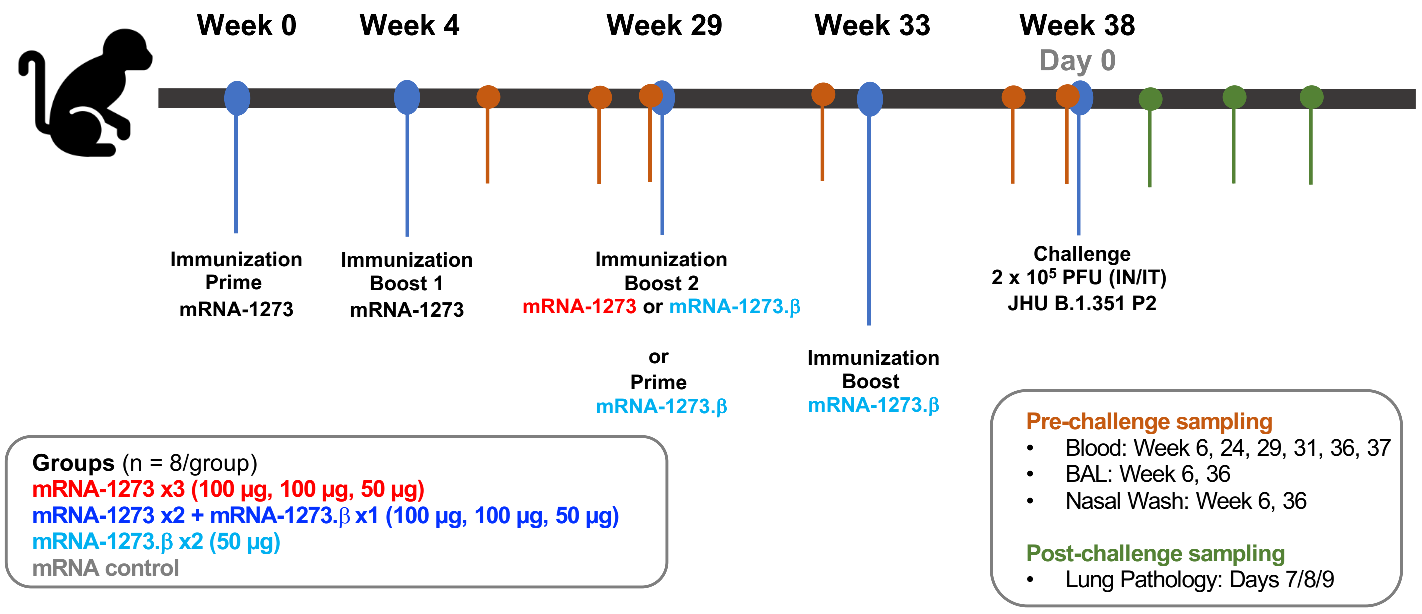


**Figure S1. Study design: Ability of mRNA-1273 booster to protect nonhuman primates (NHP) against B.1.351 (β) challenge.** Rhesus macaques (n = 8/group) were immunized on the following schedule: Group 1 (mRNA-1273 x3, red): 100 µg of mRNA-1273 at weeks 0 and 4 and 50 µg of mRNA-1273 at week 29; Group 2 (mRNA-1273 x2 + mRNA-127.β x1, dark blue): 100 µg of mRNA-1273 at weeks 0 and 4 and 50 µg of mRNA-1273.β at week 29; Group 3 (mRNA-127.β x1, light blue): 50 µg of mRNA-1273.β at weeks 29 and 33; Group 4: 100 µg of mRNA control at weeks 0 and 4. At week 38, NHP were challenged with a total of $2x10^5$ PFU of SARS-CoV-2 B.1.351 (β). The viral inoculum was administered as $3.75x10^5$ PFU in 3 mL intratracheally (IT) and $1.25x10^5$ PFU in 1 mL intranasally (IN) in a volume of 0.5 mL into each nostril. Sera were collected pre-challenge at weeks 6, 24, 29, 31 and 37. Bronchoalveolar lavages (BAL) and nasal washes were also collected at week 6 and 36. Lung pathology was assessed on days 7, 8, or 9 post-challenge.

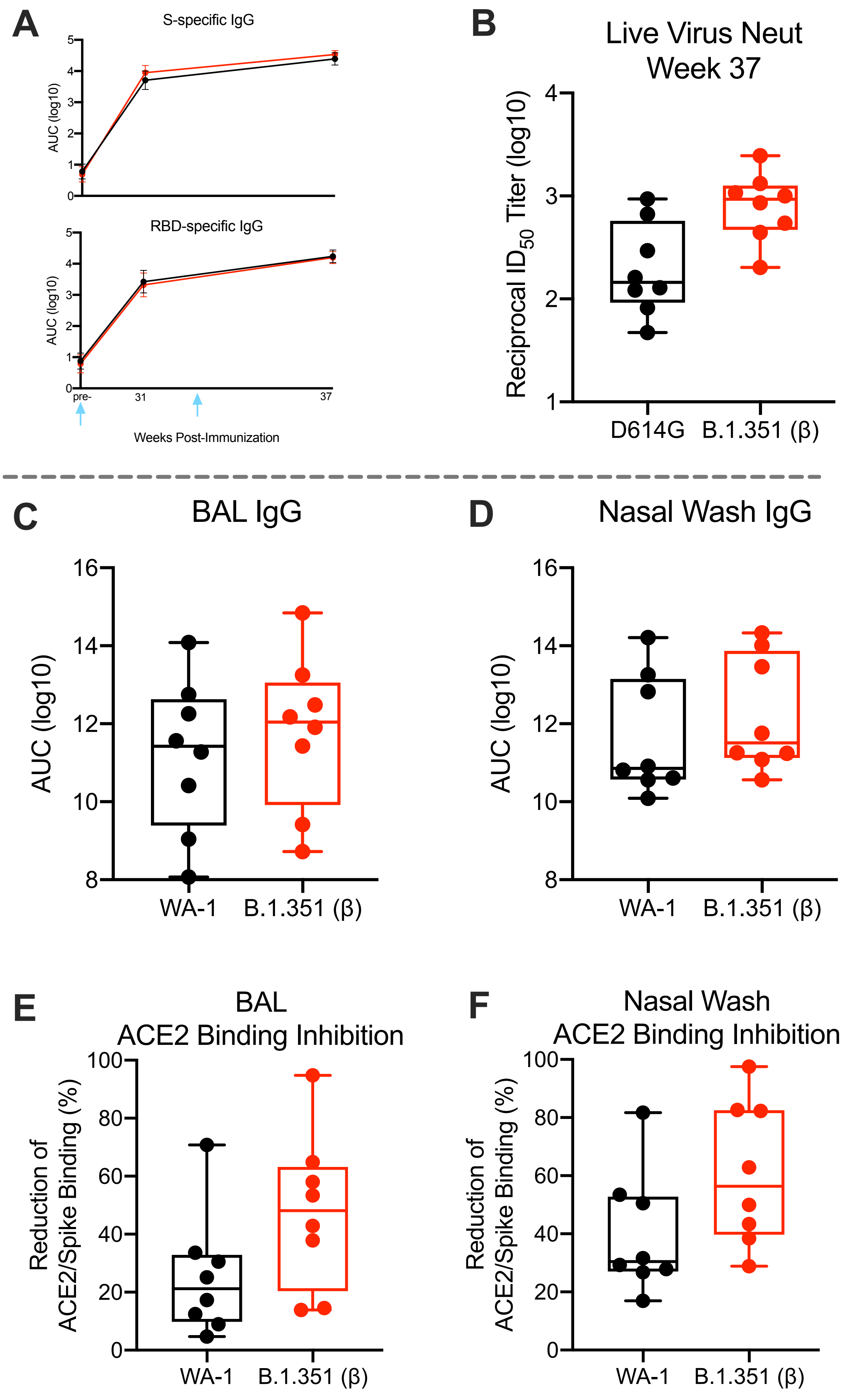
A
S-specific IgG
AUC (log10)
RBD-specific IgG
AUC (log10)
pre-
31
37
Weeks Post-Immunization
B
Live Virus Neut
Week 37
Reciprocal ID$_{50}$ Titer (log10)
D614G
B.1.351 (β)
C
BAL IgG
AUC (log10)
WA-1
B.1.351 (β)
D
Nasal Wash IgG
AUC (log10)
WA-1
B.1.351 (β)
E
BAL
ACE2 Binding Inhibition
Reduction of
ACE2/Spike Binding (%)
WA-1
B.1.351 (β)
F
Nasal Wash
ACE2 Binding Inhibition
Reduction of
ACE2/Spike Binding (%)
WA-1
B.1.351 (β)

**Figure S2.** **Antibody responses in mRNA-1273.351 control NHP.** Rhesus macaques (n = 8) were immunized with mRNA-1273.β at weeks 29 and 33, according to Figure S1. Sera collected at weeks 29, 31, and 37 and assessed for S-specific IgG to SARS-CoV-2 WA-1 (black) and β (red) by MULTI-ARRAY ELISA (A) and week 37 live virus neutralization (B). BAL (C, E) and nasal washes (D, F) collected at weeks 36 were assessed for S-specific IgG to SARS-CoV-2 WA-1 and β by MULTI-ARRAY ELISA (C-D) and inhibition of ACE2 binding to WA-1 and β S (E-F). (A) Lines represent GMT and geometric error. Arrows point to immunization timepoints. (B-E) Boxes and horizontal bars denote IQR and medians, respectively; whisker end points are equal to the maximum and minimum values. Circles represent individual NHP.

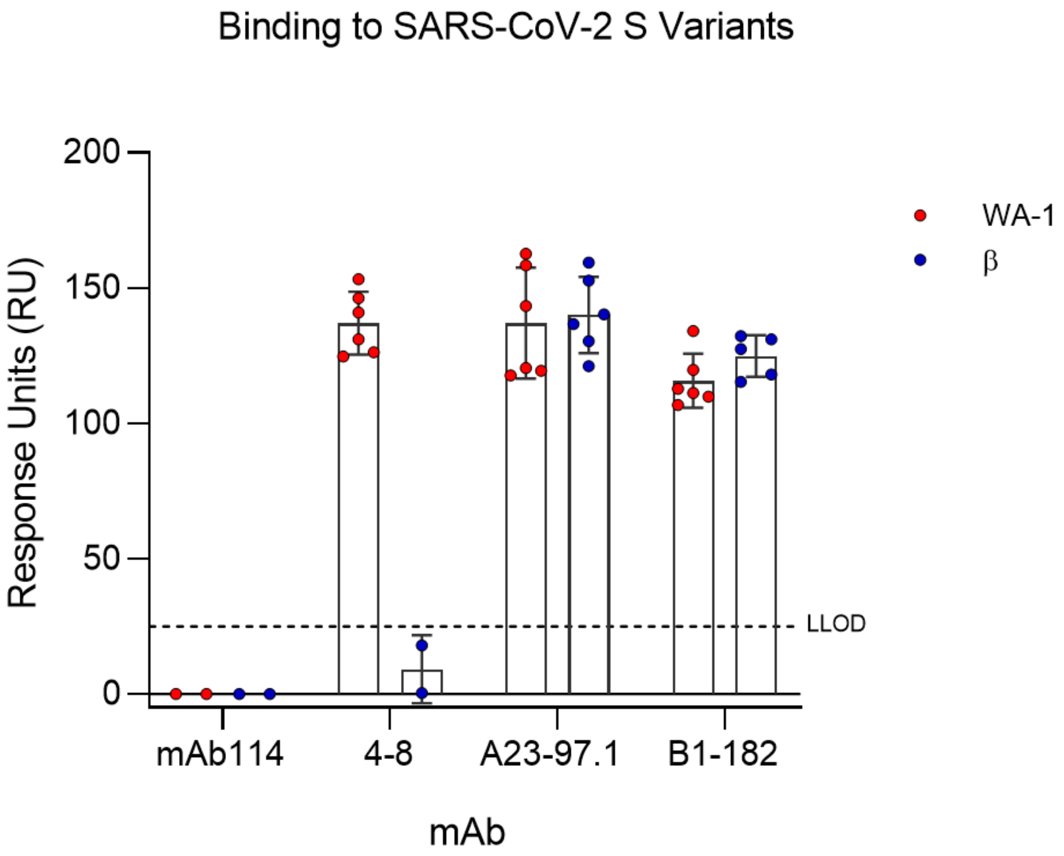


**<u>Figure S3.</u> mAb binding to SARS-CoV-2 spike variants.** Binding of mAbs to SARS-CoV-2 WA-1 and β S was evaluated using surface plasmon resonance (SPR) binding assay. Dotted line indicates assay limit of detection.

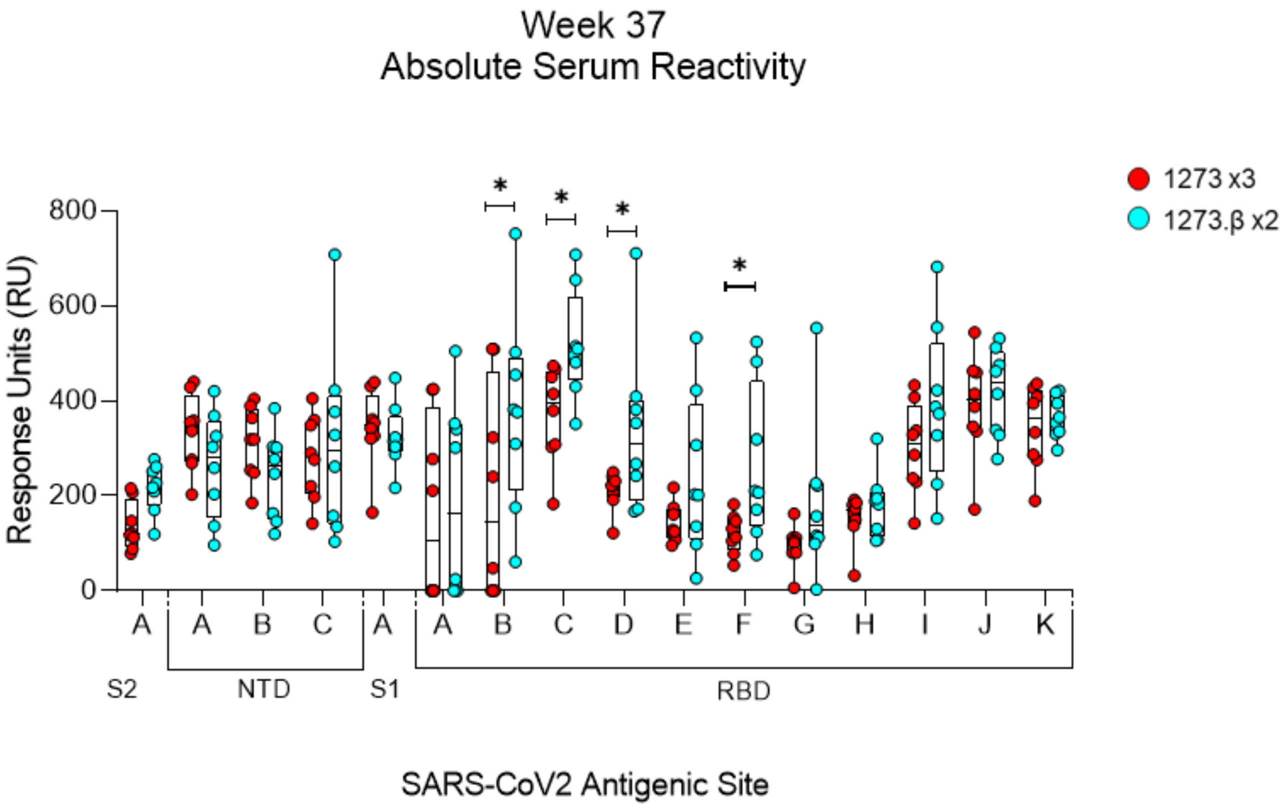


**Figure S4. Pre-challenge serum epitope profile.** Absolute serum reactivity was evaluated at Week 37 in animals receiving either mRNA-1273 x3 or mRNA-1273.β x2 vaccine regimen was determined by SPR based competition assay. Significantly ($p \leq 0.05$) increased absolute serum reactivity was observed in mRNA-1273.β x2, as compared to mRNA-1273 x3, - immunized animals to sites RBD – B, C, D, and F, represented by monoclonal antibodies CB6, A20-29.1, A19-46.1, and A19-61.1 respectively. Significant ($p \leq 0.05$) differences determined by unpaired t-test between immunization groups.

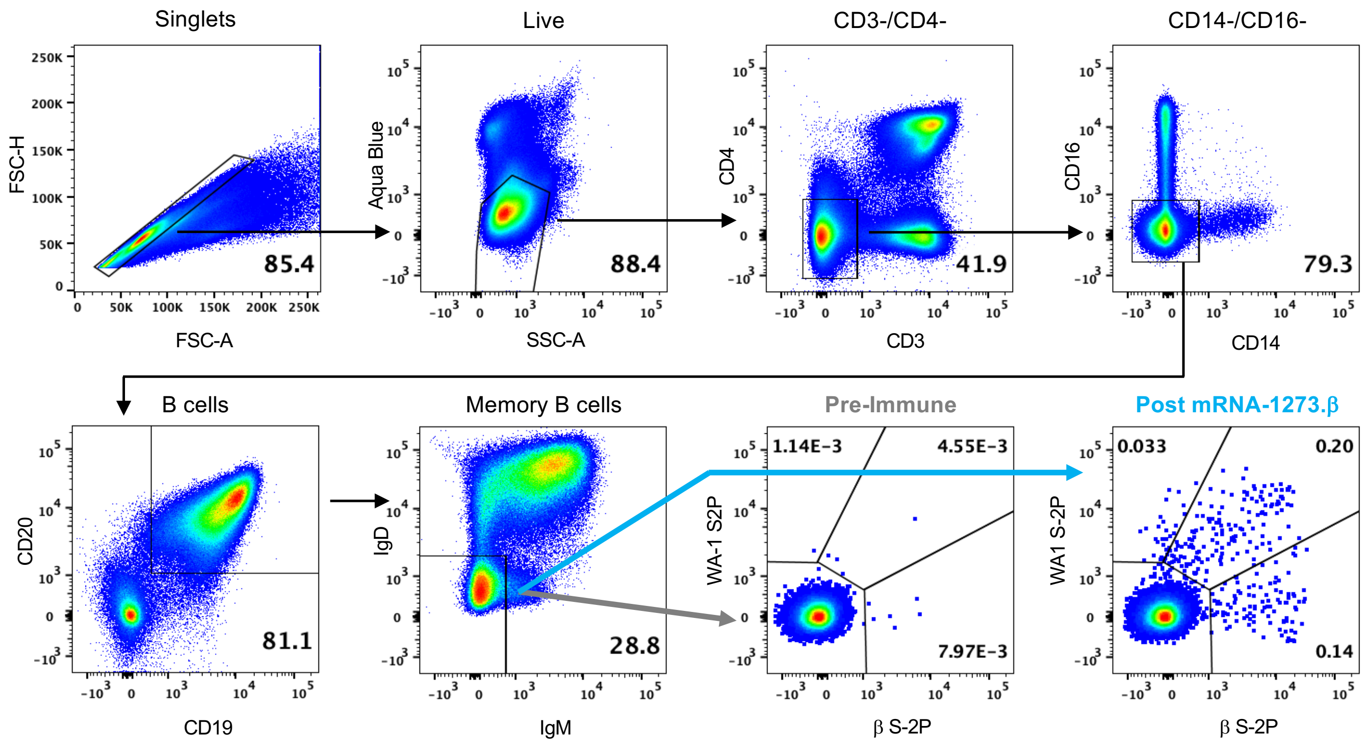


**<u>Figure S5</u>. B Cell Gating Strategy.** WA-1 S-2P, and β S-2P-specific memory B cells were identified in PBMC by gating on the following populations: singlets based on forward scatter area and height, followed by live cell gating by aqua blue and side scatter. B cells were negatively gated for CD3/CD4 and CD14/CD16, and subsequently positively gated for CD19 and CD20. Probe-specific Memory B cells were defined as IgM and IgD negative, followed by gating for WA-1 S-2P, β S-2P, and double positive populations pre-, and post immunization.

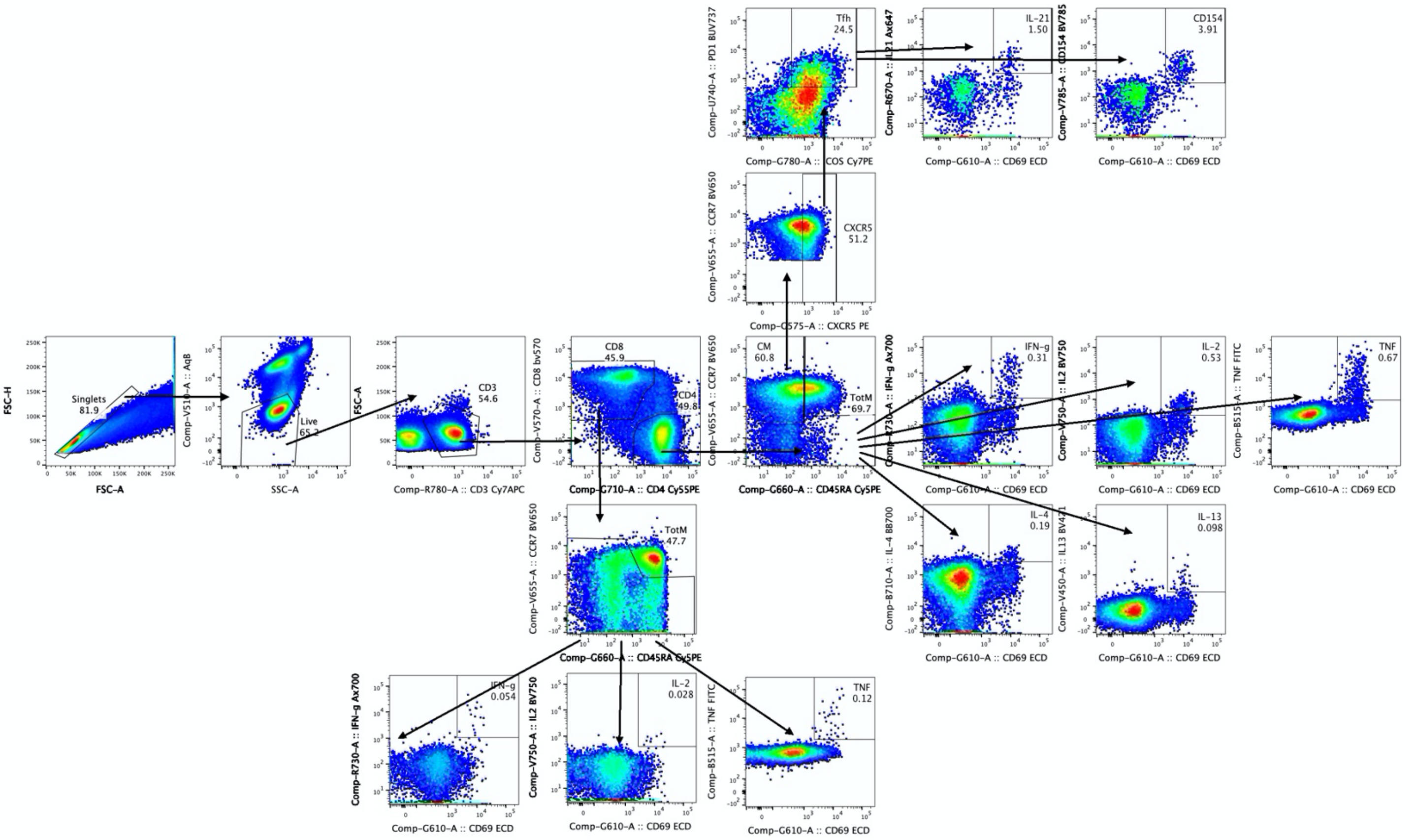


**<u>Figure S6</u>. Intracellular Staining Gating Schema.** Representative flow cytometry plots from an NHP in the mRNA-1273 x3 group at week 6. T cell populations were delineated by hierarchical gating: Singlets > Live/SSC low cells > CD3$^{+}$/FSC low cells > last CD4 and CD8 T cells. Tfh were identified by gating on central memory (CM) (CCR7$^{+}$CD45RA$^{-}$) CD4 T cells then PD-1$^{+}$ICOS$^{+}$ cells. IL-21 and CD40L (CD154) gates were then drawn on the CD69$^{+}$ Tfh population. Total memory (TotM), defined as central memory (CCR7$^{+}$CD45RA), effector memory (CCR7$^{-}$CD45RA$^{-}$), and terminal effector memory (CCR7$^{-}$CD45RA$^{+}$), CD4 and CD8 T cells were then gated. Finally, CD69$^{+}$cytokine$^{+}$ gates were drawn on total memory T cells. Th1 responses are defined as IFN-g, IL-2, or TNF positive and Th2 responses are IL-4 or IL-13 positive.

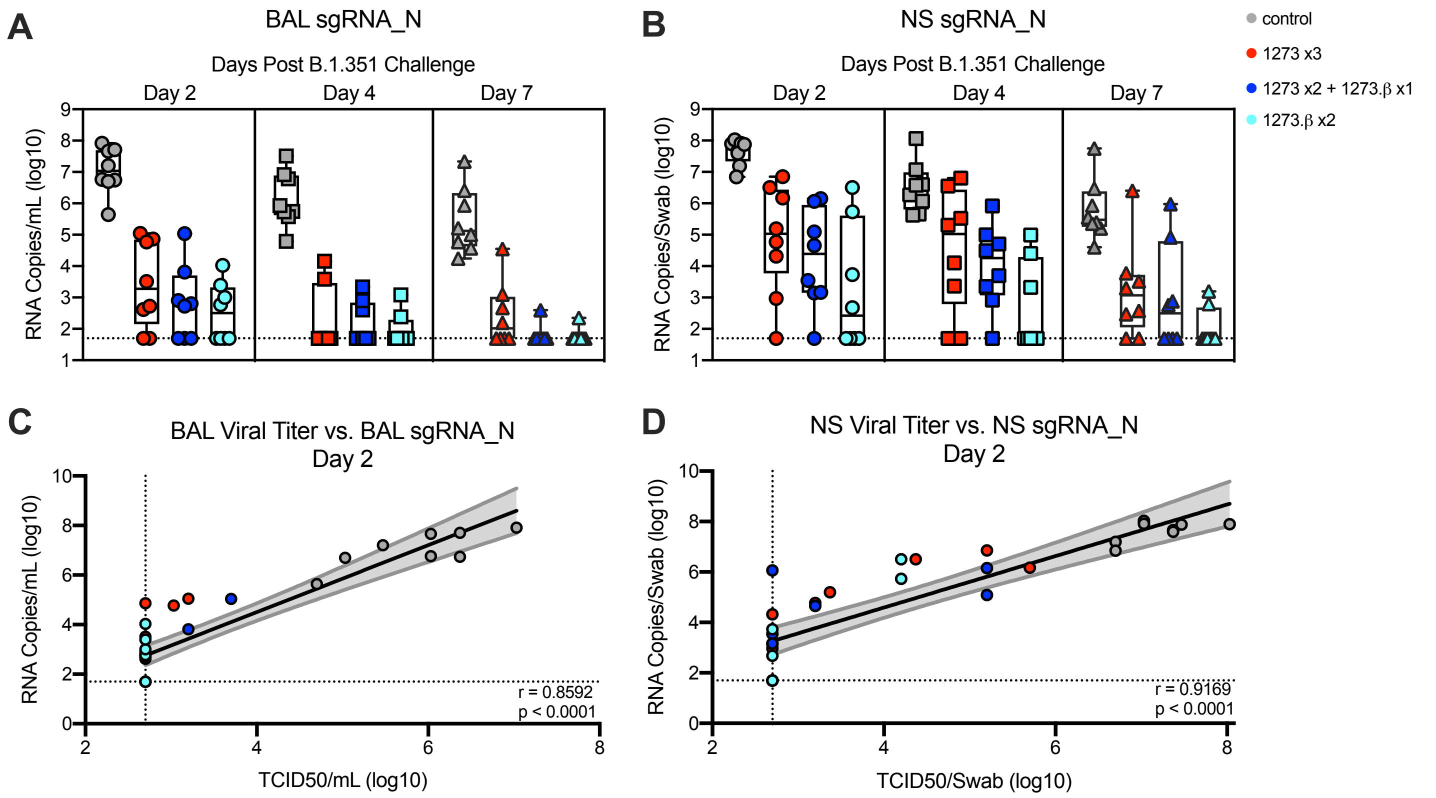


**Figure S7. Efficacy of mRNA-1273 against upper and lower respiratory B.1.351 viral replication.** Rhesus macaques were immunized and challenged as described in Figure S1. BAL (A) and nasal swabs (NS) (B) were collected on days 2 (circles), 4 (squares), and 7 (triangles) post-challenge, where applicable, and viral replication was assessed by detection of SARS-CoV-2_N sgRNA. Boxes and horizontal bars denote the IQR and medians, respectively; whisker end points are equal to the maximum and minimum values. (C-D) Plots show correlations between viral titers and sgRNA_N in BAL (C) and NS (D) 2 days post-challenge. Black and gray lines indicate linear regression and 95% confidence interval, respectively. 'r' and 'p' represent Spearman's correlation coefficients and corresponding p-values, respectively. Symbols represent individual NHP and may overlap for equal values.

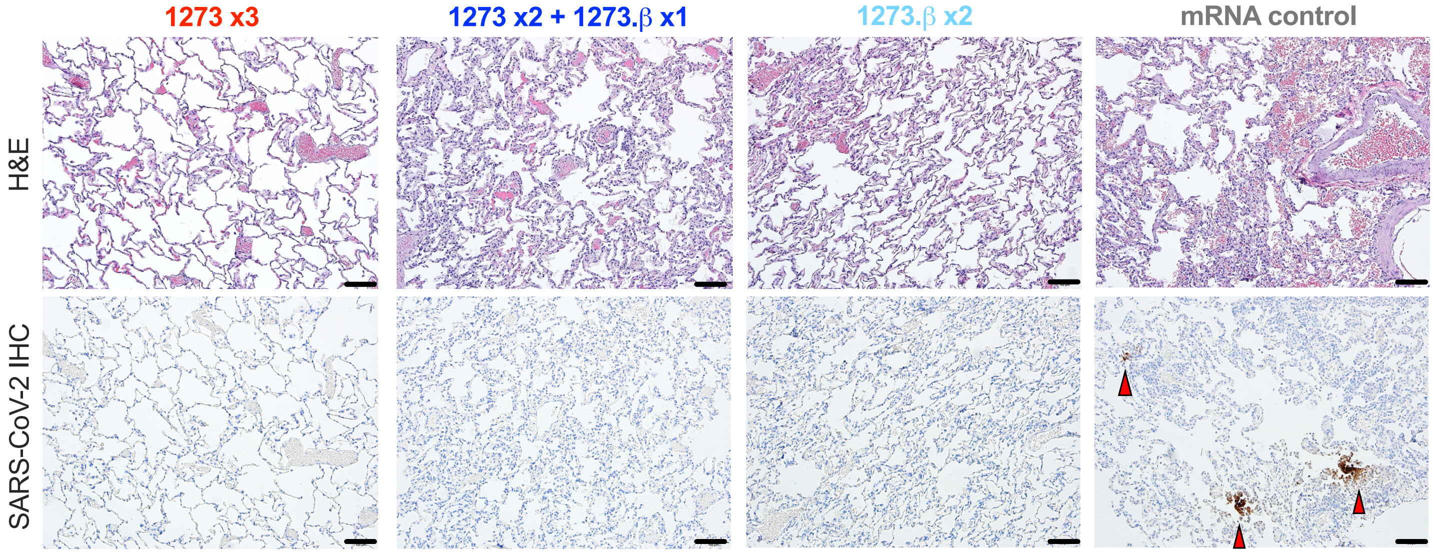


**<u>Figure S8</u>. Post-challenge lung histopathological analysis and viral detection.** Rhesus macaques were immunized and challenged as described in Figure S1. Eight days post-challenge, lung samples (n = 4/group) were evaluated for the presence of inflammation by hematoxylin and eosin (H & E) staining (left) and evidence of virus infection by immunohistochemistry (IHC) for SARS-CoV-2 viral antigen (right). Representative images show the location and distribution of SARS-CoV-2 viral antigen in serial lung tissue sections. Arrows indicate areas positive for viral antigen. Each image is taken at 10x magnification; scale bars represent 100 microns.

| Group | Animal ID | WA-1 S-specific IgG[1] (IU/mL)[2] | WA-1 RBD-specific IgG (IU/mL) |
|---|---|---|---|
| **mRNA-1273 x3** | 0NC | 369457.3 | 284665.6 |
| | 05D275 | 91364.1 | 40664.9 |
| | DGNR | 189396.4 | 132025.1 |
| | 15C225 | 345296.6 | 197959.8 |
| | DGNE | 297666.2 | 200895.6 |
| | DGTT | 174159.7 | 85585.7 |
| | DF2R | 316298.3 | 210047.0 |
| | A7V120 | 295574.2 | 173868.1 |
| | **GMT** | **239148.7** | **144489.8** |
| **mRNA-1273 x2 + mRNA-1273.β x1** | 08N027 | 280623.7 | 199878.2 |
| | 06N003 | 200510.9 | 148498.1 |
| | DGTR | 269003.2 | 198314.1 |
| | 15C108 | 173081.3 | 104116.4 |
| | 15C165 | 205204.0 | 137005.0 |
| | DF2V | 304485.3 | 185492.3 |
| | DF4G | 127770.2 | 65083.7 |
| | A7V040 | 230872.4 | 82728.2 |
| | **GMT** | **216510.0** | **130449.8** |
| **Naïve** | DH08 | *0.3*[3] | *1.6*[4] |
| | DGT0 | *0.3* | *1.6* |
| | DHGR | 44.4 | *1.6* |
| | MC44 | *0.3* | *1.6* |
| | DGD5 | *0.3* | *1.6* |
| | DF9P | *0.3* | *1.6* |
| | 14D032 | *0.3* | *1.6* |
| | HKL | *0.3* | *1.6* |
| | **GMT** | **0.6** | ***1.6*** |

**Table S1. SARS-CoV-2 WA-1 S- and RBD-specific antibody responses in international units (IU).**
[1]Sera were collected at week 37 and assessed for SARS-CoV-2 WA-1 S- and RBD-specific IgG by 4-plex MULTI-ARRAY ELISA.
[2]IgG quantities were extrapolated as IU/mL using WHO standards.
[3]S-specific IgG lower limit of detection = 0.3076 IU/mL
[4]RBD-specific IgG lower limit of detection = 1.5936 IU/mL

| | Site | mAb | |
|---|---|---|---|
| **S2** | **A** | S652-112 | |
| **NTD** | | | |
| | **A** | 4-8 | |
| | **B** | S652-118 | |
| | **C** | N3C | |
| **S1** | **A** | A20-36.1 | |
| **RBD** | | | **Barnes classification**[1] |
| | **A** | B1-182 **+** | CLASS I |
| | **B** | CB6 | |
| | **C** | A20-29.1 | |
| | **D** | A19-46.1 **+** | CLASS II |
| | **E** | LY-COV555 | |
| | **F** | A19-61.1 **+** | |
| | **G** | S309 **+** | |
| | **H** | A23-97.1 | CLASS III |
| | **I** | A19-30.1 | |
| | **J** | A23-80.1 | |
| | **K** | CR3022 | CLASS IV |

**Table S2. SARS-CoV2 Spike Antigenic Sites.** Monoclonal antibodies (mAbs) used to define antigenic sites on SARS-CoV-2 S grouped based on subdomain reactivity [S2, N-terminal domain (NTD), S1, receptor binding domain (RBD)]. Sites are defined by non-competing binding on SARS-CoV-2 S. RBD-targeted mAbs have been grouped based on their Barnes[1] classification. Monoclonal antibodies which have been shown to be neutralizing to SARS-CoV-2 β are denoted with "+".

[1]Barnes, C.O., Jette, C.A., Abernathy, M.E. *et al.* SARS-CoV-2 neutralizing antibody structures inform therapeutic strategies. *Nature* **588,** 682–687 (2020). https://doi.org/10.1038/s41586-020-2852-1

| Group | Day Post-challenge | Animal ID | Inflammation (H&E)[1] | | | IHC[2] | | |
|---|---|---|---|---|---|---|---|---|
| | | | Lc[3] | Rmid[4] | Rc[5] | Lc | Rmid | Rc |
| **mRNA-1273 x3** | 7 | 05D275 | +/- | +/- | + | - | - | +/- |
| | | DGNR | +/- | +/- | +/- | - | - | - |
| | 8 | 0NC | +/- | +/- | +/- | - | - | - |
| | | 15C225 | +/- | +/- | + | - | - | - |
| | | DGNE | +/- | + | + (cong)[6] | - | - | - |
| | 9 | DGTT | +/- | +/- | + | - | - | - |
| | | DF2R | +/- | +/- | +/- | - | - | - |
| | | A7V120 | + | +/- | + | - | - | - |
| **mRNA-1273 x2 + mRNA-1273.β x1** | 7 | 08N027 | + | +/- | + | - | - | - |
| | | DGTR | +/- | + | + | - | - | - |
| | 8 | 06N003 | +/- | + | +/- | - | - | - |
| | | 15C108 | +/- | +/- | ++ | - | - | - |
| | | 15C165 | +/- | + | + | - | - | - |
| | 9 | DF2V | + | +/- | + | - | - | - |
| | | DF4G | + (h/cong)[7] | +/- | +++ | - | - | - |
| | | A7V040 | +/- | +/- | +/- | - | - | - |
| **mRNA-1273.β x2** | 7 | DH08 | +/- | +/- | +/- | - | - | - |
| | | DGT0 | - | - | +/- | - | - | - |
| | 8 | DHGR | +/- | +/- | ++ | - | - | - |
| | | MC44 | +/- | +/- | ++ | - | - | - |
| | | DGD5 | +/+ | + | + | - | - | - |
| | 9 | DF9P | +/- | +/- | + | - | - | - |
| | | 14D032 | +/- | +/- | +/- | - | - | - |
| | | HKL | +/- | +/- | +/- | - | - | - |
| **control** | 7 | G57D | - | - | - | - | - | +/- |
| | | 15C237 | +/- | - | +/- | + | - | + |
| | 8 | DGDE | +/- | +/- | ++ | - | - | +/- |
| | | A13V200 | +/- | + | ++ | - | + | + |
| | 9 | A13V160 | +++ | +/- | +/- | + | - | - |
| | | 06N008 | +/- | ++ | +/- | +/- | + | - |

**Table S3. Assessment of lung inflammation and viral antigen on day 8 following SARS-CoV-2 B.1.351 challenge in mRNA-1273-immunized NHP.** Lung tissue, collected at days 7, 8, or 9 post-challenge, was evaluated for the presence of inflammation by H & E staining and evidence of virus infection by IHC for SARS-CoV-2 viral antigen.

[1]Inflammation scoring: - = minimal to absent, +/- = minimal to mild, + = mild to moderate, ++ = moderate to severe, +++ = severe
[2]Immunohistochemistry SARS-CoV-2 antigen (Ag): - = no detection of virus Ag, +/- = rare/occasional Ag+ foci, + = occasional - multiple Ag+ foci
[3]Lc: left caudal lung lobe
[4]Rmid: right middle lung lobe
[5]Rc: right caudal lung lobe
[6]cong: congestion
[7]h/cong: hemorrhage and congestion

| mRNA Construct ID | UNFIX-01 |
|---|---|
| Cap | C1 |
| 5’ UTR | GGGAAAUAAGAGAGAAAAGAAGAGUAAGAAGAAAUAUA<br>AGACAGCGCGUCAACAUUGCCGAAUCGCCGGGACUCAUC<br>ACAAUCUGCCUCUUGGGUUAUCUCUUGUCGGCAGAUACC<br>UUCUUGGAUCACGAAAACGCGAACAAAAUUCUUAAUCGC<br>CCGAAGCGGUAUAACUCCGGGAAACUUGAGGAGUUUCAG<br>GGCAAUCUUGAACGAGACGAGGAGAACUCCUUUGAGGAG<br>GCGAGGGAAUUUGAAAACACAGAGCGAACAACGGAGUUU<br>UGGAAGCAAUACGUAGGGGACCAGUCGAAUCCCCUCAGG<br>GGAUCUAAAGACAUCAAUAGCUACUGCCCGUUUGGGUUU<br>GAAGGGAAGAACUAGCUGACCAACAUCAAAAACGGACGC<br>UAGCAGUUUUGUAAGAACUCGGCUGACAAUAAGGUAGUC<br>UCCACAGAGGGAUACCGGCUGGCGGAGAACCAAAAAUCC<br>GAGCCCGCAGUCCCGUUCCCUUGGAGGAGCUCACAGACU<br>AGCAAGUUGACGAGAGCGGAGACUGUAUUCCCCGACGAC<br>UACGUCAACAGCACCGAAGCCGAAACAAUCCUCGAUAAC<br>AUCACGCAGAGCACUCAGUCCUUCAACUUUACGAGGGUC<br>GUAGAGGACGCGAAACCCGGUCAGUUCCCCUGGCAGGUA<br>UUGAACGGAAAAGUCGCCUUUUGAGGUUCCAUUGUCAAC<br>GAGAAGAUUGUCACAGCGGCACACUGCGUAGAAACAGGA<br>AAAAUCACGGUAGCGGGAGAGCAUAACAUUGAAGAGACA<br>GAGCACACGGAACAAAAGCGAAUCAUCAGAAUCAUUCCA<br>CACCAUAACUAUAACGCGGCAAUCAAUAAGUACAAUCAC<br>GACAUCGCACUUUUGGAGCUUGACGAACCUUUGCUUAAU<br>UCGUACGUCACCCCUAUUUGUAUUGCCGACAAAGAGUAU<br>ACAAACAUCUUCUUGAAAUUCGGCUCCGGGUACGUAUCG<br>GGCUGGGGCAGAUUCCAUAAGGGUAGAUCCGCACUGUUG<br>CAAUACCUCAGGCCCCUCGAUCGAGCCACUUGUCUGCGG<br>UCCACCAAAUUCACAAUCUACAACAAUUUCUCGGGAUUC<br>CAAGGGAGAGAUAGCUGCCAGGGAGACUCAGGGGGUCCC<br>CACACGGAAGUCGAGGGGACGUCAUUUCUGACGGGAAUU<br>AUCUCGGGAGAGGCGAAGGGGAACAUCUACACUAAAUCA<br>CGGUUCAAUUGGAUCAAGGAAAAGACGAAACUCACGUGA<br>UAAUAGGCUGGAGCCUCGGUGGCC |
| ORF of mRNA Construct | AUGCUUCUUGCCCCUUGGGCCUCCCCCCAGCCCCUCCUCC<br>CCUUCCUGCACCCGUACCCCCGUGGUCUU |
| 3’ UTR | UGAAUAAAGUCUGAGUGGGCGGC |
| Corresponding amino acid sequence | MLLAPWASPQPLLPFLHPYPRGL |
| PolyA tail | 100 nt |

**Table S4. Sequence of control mRNA.**

## A

| Variant | MUTLI-ARRAY ELISA |
|---|---|
| **WA-1** | *No mutations* |
| **D614G** | *N/A*[1] |
| **B.1.1.7** | Δ69-70, Δ144, 501Y, 570D, 614G, 681H, 716I, 982A, 1118H |
| **P.1** | 18F, 20N, 26S, 138Y, 190S, 417T, 484K, 501Y, 614G, 655Y, 1027I, 1176F |
| **B.1.351** | 18F, 80A, 215G, Δ242-244, 246I, 417N, 484K, 501Y, 614G,701V |

## B

| Variant | Pseudovirus | |
|---|---|---|
| | **Lentiviral** | **VSV** |
| **WA-1** | *N/A* | *N/A* |
| **D614G** | 614G | 614G |
| **P.1** | 18F, 20N, 26S, 138Y, 190S, 417T, 484K, 501Y, 614G, 655Y, 1027I, 1176F | *N/A* |
| **B.1.351** | 18F, 80A, 215G, Δ242-244, 246I, 417N, 484K, 501Y, 614G,701V | 18F, 80A, 215G, Δ242-244, 246I, 417N, 484K, 501Y, 614G, 701V |
| **B.1.429** | S13I-W152C-L452R-D614G | *N/A* |
| **B.1.526** | L5F-T95I-D253G-E484K-D614G-A701V | *N/A* |
| **B.1.617.2** | T19R, G142D, del156-157, R158G, L452R, T478K, D614G, P681R, D950N | *N/A* |

## C

| Variant | Live Virus |
|---|---|
| **WA-1** | *N/A* |
| **D614G** | 614G |
| **B.1.351** | 18F, 80A, 215G, Δ242-244, 417N, 484K, 501Y, 614G, 701V |

**Table S5. Spike mutations included in Meso-scale ELISA (A), pseudovirus (B) and live virus (C) assay reagents as compared to Wuhan-1 strain (Genbank #: MN908947.3).**

[1]N/A: not applicable
